# Spatiotemporal cellular landscape of the human utricle sensory epithelium

**DOI:** 10.64898/2026.06.25.734616

**Authors:** Weisheng Liang, Ryosuke Yamamoto, Emilia Luca, Alain Dabdoub

## Abstract

Vestibular dysfunction affects individuals of all ages and becomes increasingly common with age. Despite the essential role of the utricle in balance, its molecular states and tissue architecture in humans remain poorly defined. Here, we integrated paired single-nucleus transcriptomic and epigenomic profiling with imaging-based spatial transcriptomics to characterize the human fetal utricle and its spatiotemporal patterning. We resolved transcriptionally heterogeneous, spatially segregated populations of sensory and nonsensory epithelial cells and reconstructed sensory cell differentiation across three gestational ages, showing that these cells acquire region-specific transcriptional signatures before final subtype specification. We further uncover a progressive decline in nonsensory cell proliferation accompanied by chromatin remodelling, as well as transitional epithelial populations with distinct spatial and regulatory programs. Together, these data define the molecular and spatial dynamics of the human fetal utricle and reveal cell states and regulatory pathways that provide a foundation for studying vestibular disorders and regeneration.

**HIGHLIGHTS:**

- Spatial identity of utricular sensory hair cells precedes full subtype specification
- Distinct transcriptional regulators control hair cell fate and regional patterning
- Supporting cells remodel chromatin as their proliferative capacity declines
- Transitional epithelial cells adopt ordered states along the utricular border
- Sensory and nonsensory cells show divergent regulatory and signalling programs

## INTRODUCTION

The vestibular system of the inner ear detects head orientation and motion to maintain balance. Within this system, the utricle senses linear acceleration and head tilt on the horizontal plane via mechanosensory hair cells present in its sensory epithelium. Damage or loss of vestibular hair cells occurs with aging, noise exposure, and ototoxic therapeutics, including aminoglycoside antibiotics and cisplatin^1–4^ leading to dizziness, vertigo, and an increased risk of falls, a leading cause of serious injury. The prevalence of balance dysfunction increases with age, affecting over 5% of children and adolescents, nearly 40% of adults, and up to 50% in the elderly, thereby imposing a major healthcare burden.^5–8^ Because mature mammalian vestibular hair cells have limited regenerative capacity,^9–12^ their loss can lead to progressive and persistent dysfunction, underscoring the need to define developmental cell states and regulatory programs that might be harnessed for vestibular repair and balance restoration.

During utricle development, prosensory progenitor cells give rise to both hair cells and the surrounding supporting cells, which maintain structural integrity and ion homeostasis within the sensory epithelium.^13,14^ In mammals, supporting cells persist after hair cell loss and can regenerate hair cells during early developmental stages, but this capacity declines sharply with maturation. As the utricle matures, the sensory epithelium diversifies into distinct hair cell and supporting cell subtypes and becomes spatially organized into a central striolar region and a peripheral extrastriolar region. The sensory epithelium is additionally bordered by transitional epithelial cells, a nonsensory population that forms an interface with the epithelial roof. Understanding when and how these cellular and regional identities emerge is likely to reveal mechanisms that support hair cell regeneration and inform efforts to restore function to mature vestibular organs.

Recent transcriptomic and epigenomic studies in mice have begun to characterize dynamic changes that accompany utricle development and maturation. In the mouse utricle, hair cell formation and maturation are robust during late embryonic and neonatal stages, with hair cell numbers doubling in the first postnatal week.^15^ In contrast, available histological studies suggest that human utricle hair cells appear morphologically mature and stabilize their number by gestational week (W) 16.^16,17^ Although single-cell transcriptomic profiling has revealed heterogeneity in the adult human utricle, prior datasets have largely represented older adults and disease-associated contexts^10,18,19^ and the developing fetal utricle has yet to be characterized at comparable resolution. It remains unclear how human utricular cell states change across gestation, when spatial domains including striolar and extrastriolar regions, are established, and which regulatory programs underlie these transitions. Addressing these gaps is essential for understanding the spatiotemporal basis of utricle function and for identifying cell populations with regenerative potential.

Here, we present a multidimensional analysis of the developing human utricle sensory epithelium using single-nucleus multiomic profiling across gestational ages W12, W15, and W19, integrated with spatial transcriptomics to validate and map gene expression within intact tissue. This gestational window captures a period when the utricle sensory epithelium is undergoing molecular and regulatory maturation, including hair cell specification and hair cell bundle formation. This allowed us to define hair cell populations and their molecular dynamics during development. We further resolved regionally patterned supporting cells and uncovered four spatially distinct transitional epithelial cell subpopulations. Collectively, these integrated multimodal data link developmental cell states, regulatory programs, and spatial organization in the human fetal utricle at an unprecedented resolution, highlighting a period during which the sensory epithelium remains transcriptionally and epigenetically dynamic and establishing a foundation for interrogating vestibular disorders and regenerative strategies.

## RESULTS

### Transcriptomic analysis defines the cellular composition of the fetal utricle

Utricles were dissected from W15 and W19 bony labyrinths (n=2 each; Figure 1A). Representative immunostaining for F-actin (hair bundles), MYO7A (hair cells), and SOX2 (supporting cells and developing hair cells) showed the mosaic organization of the utricle sensory epithelium (Figure 1B). Sensory epithelia were delaminated and processed for single-nucleus multiomic sequencing to generate paired transcriptomic and chromatin accessibility profiles from the same nucleus (Figure 1C). After performing quality controls, 12,761 nuclei were retained for downstream analysis with Seurat^20^ with a median of 2,198 genes per nucleus (Figures 1D and E; Table S1a). Transcriptomic integration across ages using reciprocal principal component analysis (RPCA), followed by Leiden clustering, resolved three major epithelial cell populations confirmed by *EPCAM* enrichment across the dataset (Figure 1F). Cell types were annotated based on established markers for hair cells (*OTOF*), supporting cells (*SOX2*; also enriched in a subset of hair cells), and transitional epithelial cells (*CNMD*).

**Figure 1:**
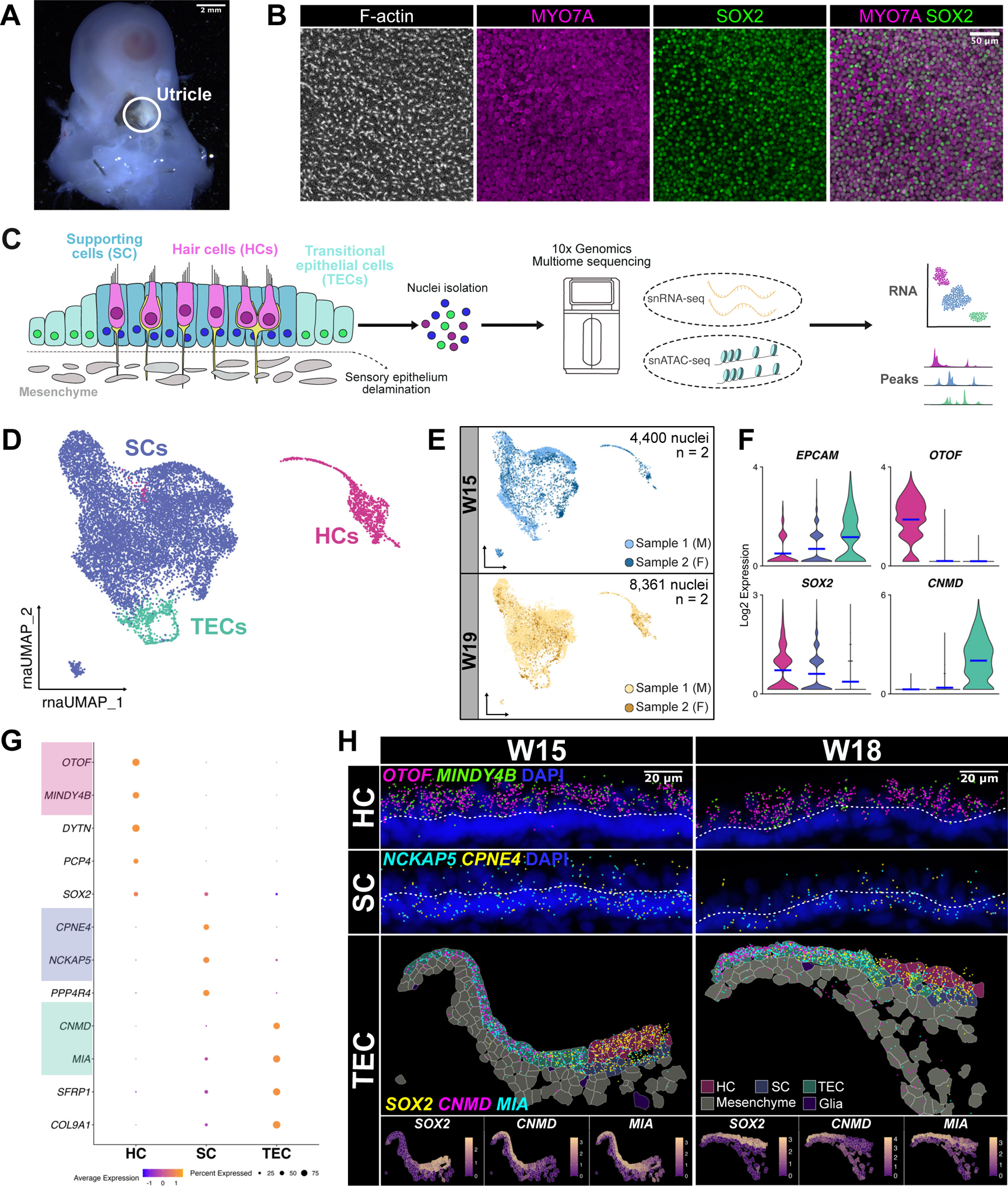
Cell types of the human fetal utricle. (A) W16 bony labyrinth with the utricle exposed. Scale bar: 2 mm. (B) Whole-mount immunohistochemical staining of a W16 utricle showing stereocilia bundles (F-actin), hair cells (MYO7A), supporting cells (SOX2^+^ MYO7A^-^), and merged MYO7A and SOX2 staining. Scale bar: 50 µm. (C) Schematic of the single-nucleus multiome workflow from sensory epithelium dissection through sample processing and downstream statistical analysis. (D) UMAP projection of W15 and W19 snRNA-seq data showing annotation of hair cells (HCs), supporting cells (SCs), and transitional epithelial cells (TECs). (E) Nuclei distribution across ages and samples (n=2 per stage). (F) Violin plots of known sensory epithelium markers: *EPCAM* for epithelial cells, *OTOF* for hair cells, *SOX2* for supporting cells with expression in a subset of hair cells, and *CNMD* for transitional epithelial cells. (G) Dot plot of cell type markers identified through differential gene expression analysis. Genes validated in (H) are highlighted. (H) *In situ* validation of cell type-specific markers. Hair cells: *MINDY4B*; supporting cells: *NCKAP5* and *CPNE4*. The dashed line separates hair cell and supporting cell nuclei within the same tissue segment. Scale bar: 20 µm. Spatial feature plots show expression of transitional epithelial cell genes *CNMD* and *MIA* alongside *SOX*2, n=3.

We further distinguished these three populations with differential gene expression analysis. We identified 2,424 differentially expressed genes (DEGs) in hair cells, 338 in supporting cells, and 410 in transitional epithelial cells (non-parametric Wilcoxon rank-sum test, *P_adj_* < 0.05 and log_2_FC ≥ 2, Table S2a), with representative genes highlighted (Figure 1G). To assess disease relevance, we examined genes associated with vestibular dysfunction and hearing loss^19^ using a gene detection threshold of expression in ≥ 25% of cells in any cell type. We detected 56% (38/68) of vestibular impairment-associated genes, and 49% (44/90) of hearing loss-associated genes in both hair cells and nonsensory cells (Figures S1A and B; Table S1b and c). Thus, our fetal utricle datasets capture broad expression of genes linked to inner ear disorders, underscoring their clinical relevance for investigating human vestibular dysfunction.

To examine gene expression within the intact tissue architecture, we designed a custom gene panel informed by our multiomic analysis (Table S3) and utilized the 10x Genomics Xenium spatial transcriptomics platform to profile W15 and W18 utricle cross-sections (n=3 each), with the latter used for spatial support of the W19 dataset. Clustering and annotation of Xenium cells using canonical markers identified hair cells (*OTOF*^+^), supporting cells (*SOX2*^+^), transitional epithelial cells (*CNMD^+^*), epithelial roof cells (*CITED1*^+^), mesenchyme (*EPCAM*^-^), and glia (*EPCAM*^-^*SOX2*^+^; Figures S1C-H). Visualization of individual transcripts (Xenium Explorer, v.3.2.0) confirmed spatially restricted expression of the human-specific hair cell marker *MINDY4B* as previously described,^18^ as well as supporting cell genes *NCKAP5* and *CPNE4* (Figures 1G and H). Hair cell and supporting cell markers localized to distinct nuclear layers within the sensory epithelium, consistent with their expected organization. Supporting cell transcripts detected in the hair cell nuclei layer reflect cytoplasmic interdigitation around hair cells. In contrast, the transitional epithelial markers *CNMD* and *MIA* increased progressively outside the sensory region, while *SOX2* decreased in a reciprocal pattern (Figure 1H). Together, these transcriptomic and spatial analyses define the major cell populations of the fetal utricle and reveal spatially organized epithelial compartments.

### Hair cell transcriptional states resolve differentiation and spatial identity

To define hair cell differentiation across development, we extracted hair cells from each age group separately and performed independent subclustering. At W15, Leiden clustering resolved three hair cell subpopulations. Most W15 hair cells expressed *SOX2*, consistent with an early differentiation stage.^21,22^ One cluster was enriched for the immature hair cell marker *ATOH1*, the master regulator for hair cell formation; another was enriched for the more differentiated hair cell marker *TMC2*, which encodes a mechanotransduction channel subunit; and a third cluster enriched in *CHRNG* was positioned between these two states, (Figures 2A and B). Pseudotime analysis using Monocle3^23^ identified changing expression of hair cell differentiation genes (*ATOH1*, *HES6, IGFBP3*) and stereocilia-associated genes (*MYO3B, PTPRQ, XIRP2*) across these clusters, which indicated that the *CHRNG*^+^ cluster likely represents an intermediate transitional state between immature and more mature hair cells (Figures 2C and S2A; Table S4a). These data characterized transcriptional trajectories of W15 hair cells consistent with stepwise differentiation from an *ATOH1^+^* immature state toward more mature *TMC2*^+^ states.

**Figure 2:**
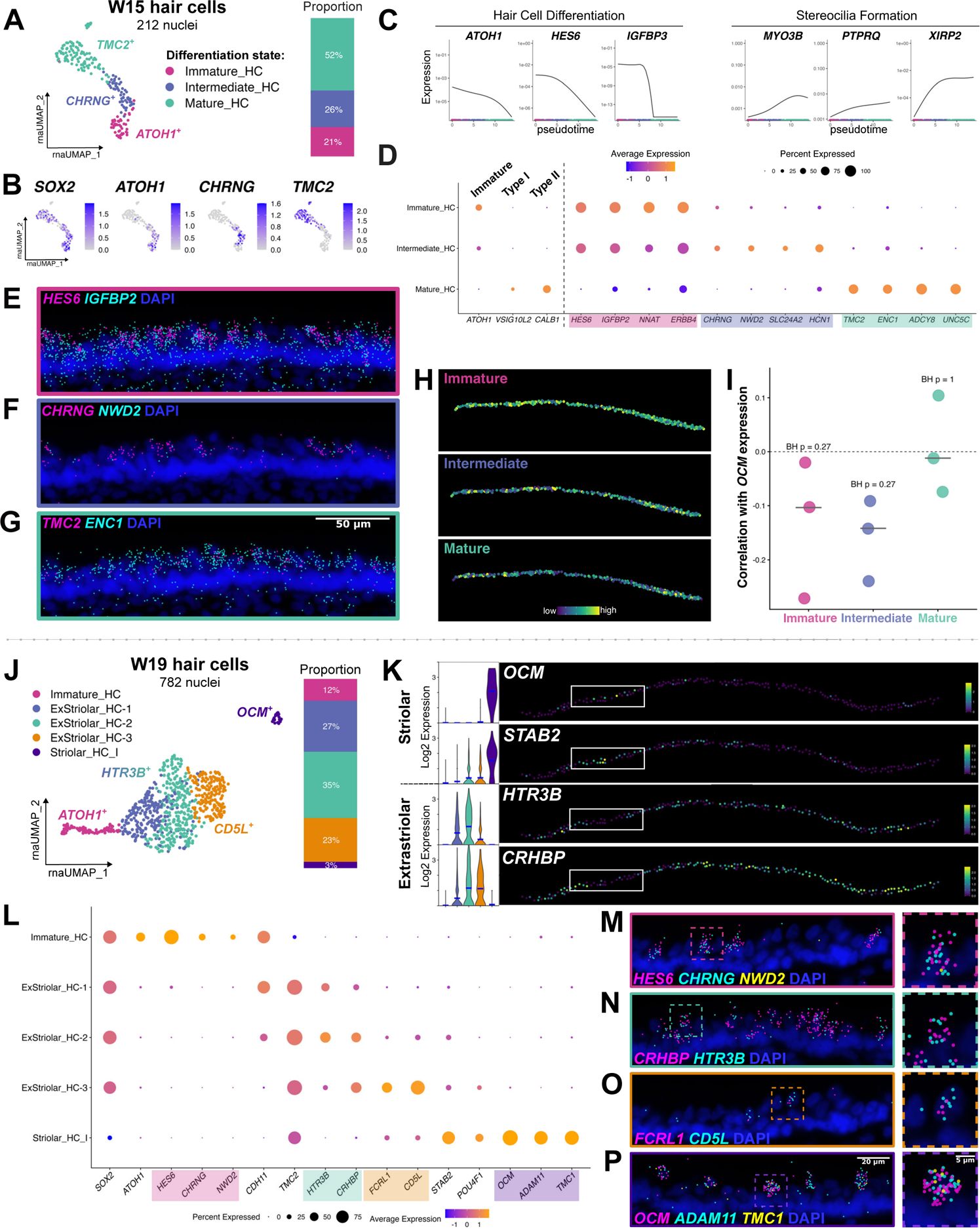
Transcriptomic and spatial identities of utricle hair cells at W15 and W19. (A) UMAP projection and cluster proportions of W15 hair cells, n=2. (B) Feature plots showing *SOX2*, *ATOH1, CHRNG*, and *TMC2* at W15. (C) Expression of early hair cell differentiation and stereocilia formation genes across pseudotime. (D) Dot plot of hair cell subtype markers and enriched genes in each W15 hair cell cluster. (E-G) *In situ* validation of genes enriched in immature (E), intermediate (F), and mature (G) hair cell clusters, n=3. Scale bar: 50 µm. (H) Spatial plot of module scores for each hair cell cluster (n=3). (I) Spearman correlations between *OCM* expression and W15 maturation state module scores. Points represent W15 Xenium samples; bars indicate the median. BH-adjusted *P*-values test whether correlations differ from 0. (J) UMAP projection and cluster proportions of W19 hair cells, n=2. (K) Violin plots and spatial feature plots of regionally enriched genes in W19 hair cells. Striolar: *OCM*, *STAB2*; Extrastriolar: *HTR3B*, *CRHBP*, n=3. (L) Dot plot of genes enriched in W19 hair cell clusters. (M-P) *In situ* validation of genes highlighted in (L), n=3. Scale bars: 20 µm, 5 µm for insets.

Differential expression analysis identified genes enriched in each W15 hair cell differentiation state (Figure 2D; *P_adj_* < 0.05 and log_2_FC ≥ 1, Table S2b), with representative markers validated by *in situ* to be expressed throughout the hair cell layer. (Figures 2E-G, S2B). The intermediate population, defined by *CHRNG* which encodes the gamma subunit of the acetylcholine receptor, also expressed *NWD2*, implicated in cholinergic signal transduction.^24^ Although *NWD2* has not been characterized in the inner ear, *Chrng* expression has been reported in mouse vestibular hair cells during embryonic and early postnatal stages.^25^ This may reflect a developmental period where hair cells acquire cholinergic signalling features, which occurs during development and persists in vestibular type II hair cells.^26,27^

We next assessed whether type I and type II hair cell identities are transcriptionally resolved at W15. Established adult human markers for type I (*VSIG10L2* and *OCM*) and type II hair cells (*CALB1*)^10,18,19^ are expressed in *TMC2*^+^ hair cells (Figure 2D). Although only represented by a few cells, *OCM* marked a discrete subset of hair cells that separated in UMAP space, and localized centrally within the sensory epithelium (Figure S2C), corresponding to the striolar region of the utricle and consistent with the spatial pattern of *Ocm* in mice.^28,29^ In contrast, *CALB1* was expressed across *TMC2*^+^ hair cells while *VSIG10L2* was detected in a subset (Figure S2C); thus, the striolar type I identity is spatially defined by W15.

To test whether regional identity was linked to hair cell differentiation at W15, we calculated module scores for immature-, intermediate-, and mature hair cell programs in Xenium hair cells and correlated these scores with *OCM* expression as a proxy for striolar identity (Table 1d). *OCM* expression was not significantly associated with any differentiation state-module score (Figure 2I), suggesting that regional identity and hair cell differentiation are not coupled at this stage.

To determine how hair cell differentiation progresses with age, we examined hair cell states in the W19 utricle. Subclustering resolved five populations, with *TMC2^+^* hair cells now predominant (Figure 2J and S2D). However, retention of *SOX2* in most *TMC2*^+^ hair cells suggests that they are incompletely specified, developmentally preceding the emergence of *SOX2-*negative type I hair cells. *VSIG10L2*^+^ and *CALB1*^+^ cells also demonstrated limited separation into distinct type I and type II clusters (Figure S2D). Spatial expression in W18 was comparable to W15 cross-sections, with *OCM* representing the striolar region while *VSIG10L2* and *CALB1* displayed more widespread expression (Figure S2E), and over 60% of hair cells co-expressed both genes (Table S1f). Using top type I and type II hair cell markers from adult utricle datasets (Table S1e),^10,18^ we calculated module scores for each W19 hair cell cluster and found that the striolar cluster scored high for type I identity (Figure S2F). ExStriolar_HC-1 and HC-2 displayed marginally higher type II scores, while ExStriolar_HC-3 was approximately equivalent between both. These findings demonstrate that the striolar type I identity is established before distinct transcriptional specification of hair cell subtypes.

Among the mature hair cell populations, extrastriolar hair cells were enriched for genes including *HTR3B*, *CRHBP*, *WDR49*, *SEMA3E*, and *CD5L,* along with *CIB3* as a known extrastriolar marker.^30^ Striolar type I hair cells expressed genes including *STAB2*, *DGKI*, *ADAM11*, *TMC1*, and *TAC1* (Figures 2K and S2G). Notably, *Tmc1* was previously reported in extrastriolar hair cells in mice,^31^ indicating species-specific differences in the regionalization of this mechanotransduction channel. Differential expression analysis identified additional markers distinguishing immature, extrastriolar, and striolar type I hair cell populations, which we validated by *in situ* using representative genes from each state (Figures 2L-P; *P_adj_* < 0.05 and log_2_FC ≥ 1; Table S2c). The most immature population, marked by *HES6*, also retained *CHRNG* and *NWD2* expression, resembling the intermediate differentiation state observed at W15. Among the extrastriolar populations, ExStriolar_HC-1 and ExStriolar_HC-2 had higher expression of *HTR3B,* which encodes a ligand-gated ion channel involved in fast synaptic transmission and has been detected in the adult mouse vestibular ganglion.^32^ Extrastriolar_HC-3 specifically expressed FCRL genes (*FCRL1*/*2*) and *CD5L*, associated with immune-related functions. We did not observe a spatial difference in the expression of these cluster-enriched genes, suggesting that these three extrastriolar clusters may represent different maturation and specification states. Together, these analyses delineate transcriptional programs associated with states of vestibular hair cell differentiation and spatial identity.

### Trajectory analysis reconstructs hair cell maturation across gestational ages

To define changing hair cell states across development along a broader maturation continuum, we generated a W12 dataset representing an earlier developmental state. W12 right and left utricles (n = 1) were pooled for snMultiome sequencing, yielding 4,370 nuclei (Figures S3A and B). We retained only epithelial nuclei (*EPCAM*^+^) for analysis (Figures S3C and D). Subclustering of W12 hair cells resolved five subpopulations (Figures S3E, Table S2d). Three *ATOH1*^+^ clusters were identified, including a *NOTCH1*^+^/*LGR5*^+^ Notch pathway-associated population, a *DLL3*^+^ population with detectable *HES1/HES5* expression that did not reach statistical significance, consistent with lateral inhibition programs described in developing cochlear hair cells, and an *NWD2*^+^ population represented an intermediate state (Figure S3F).^33^ These features suggest that W12 captures a nascent population of hair cells along with immature and intermediate differentiation states. Two *TMC2*^+^ clusters expressed markers of striolar (*STAB2*^+^) and extrastriolar (*HTR3B*^+^) hair cells (Figure S3F). To relate hair cell states across gestational ages and infer their developmental trajectories, we integrated hair cells from W12, W15, and W19 using RPCA and generated a dataset of 1,606 nuclei. Unsupervised clustering resolved seven populations spanning distinct differentiation states and spatial identities, which we annotated using marker genes defined at W15 and W19 (Figures S3G and H; Table S2e). A cluster enriched in ribosomal and mitochondrial genes was excluded from downstream trajectory analysis (Figures S3I), and the remaining hair cells were re-normalized prior to pseudotime inference (Figures 3A; Table S2f). Within the integrated dataset, the nascent *HES1^+^* hair cell cluster was derived exclusively from W12 (Figure S3G). Immature and intermediate populations (*ATOH1*^+^/*HES6*^+^; *CHRNG*^+^/*NWD2*^+^) decreased proportionally with advancing gestational age, whereas *TMC2^+^* populations became increasingly represented (Figures 3B and C). The relative abundance of extrastriolar hair cells also increased while striolar hair cells were similarly represented across gestational age (Figure 3B). Representative cluster-enriched genes are summarized in Figure 3D.

**Figure 3:**
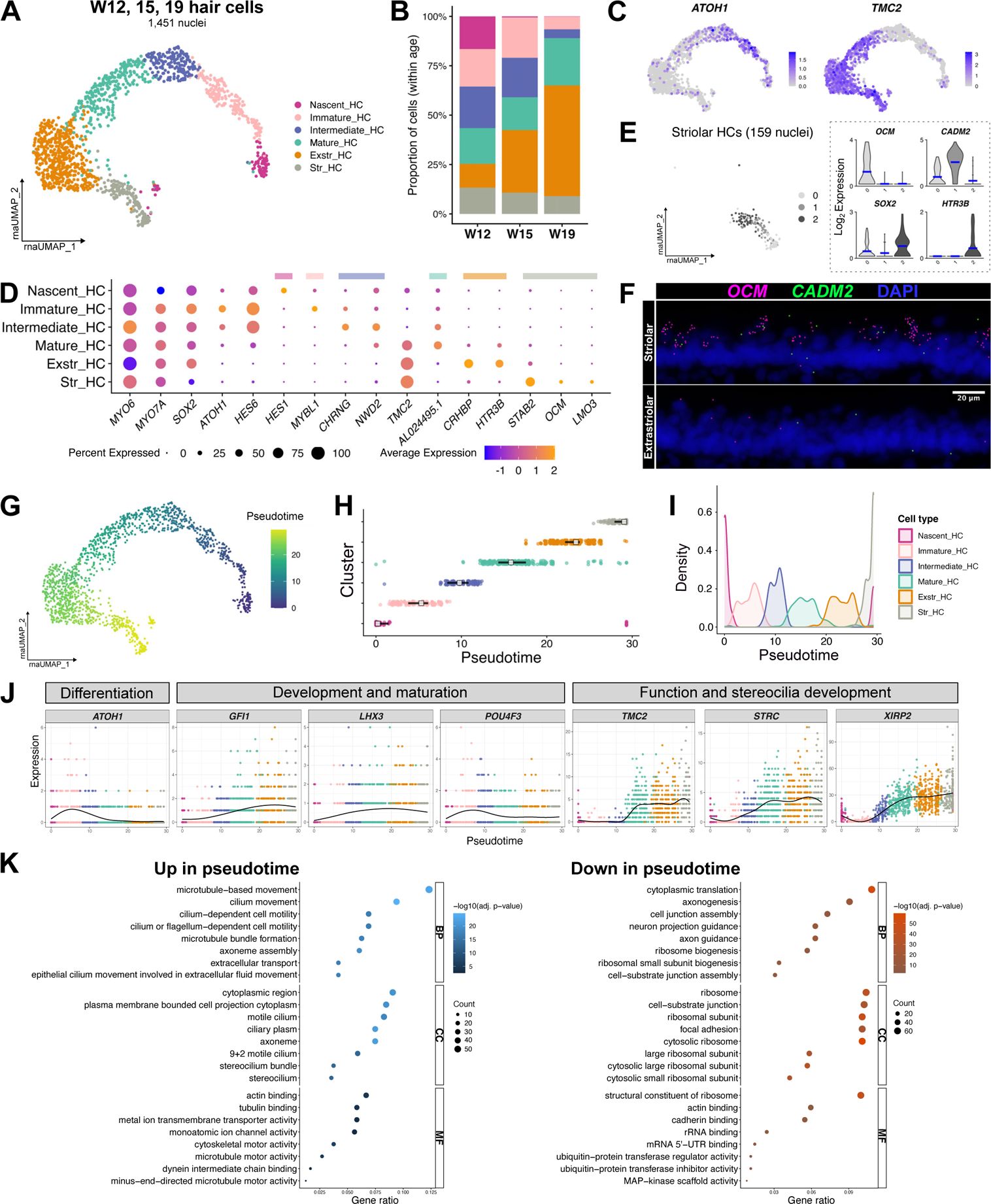
Hair cell development trajectory from W12 to W19. (A) Transcriptional UMAP projection of integrated hair cells from W12 (n=1), W15 (n=2), and W19 (n=3) datasets. (B) Proportion of hair cells belonging to each integrated cluster within each stage. (C) Feature plots of *ATOH1* and *TMC2* showing immature and mature hair cell populations. (D) Dot plot of the top differentially expressed genes in each cluster. (E) Subclustering of striolar hair cells. Inset shows violin plots of cluster-enriched genes. (F) Expression of *OCM* and *CADM2* in striolar and extrastriolar regions, n=3. (G) UMAP projection of integrated hair cells coloured by pseudotime, inferred by Monocle3. (H) Distribution of pseudotime values by cluster. White squares indicate median pseudotime; black lines indicate interquartile range. (I) Density distribution of cells from each cluster across pseudotime. (J) Expression of genes inferred to change across pseudotime, grouped into three categories. Dots represent gene expression in individual cells coloured by cluster; black lines indicate the smoothed expression trend across pseudotime. (K) GO terms for biological process (BP), cellular component (CC), and molecular function (MF) enriched among genes upregulated or downregulated across pseudotime.

Because *OCM^+^* hair cells represent only a subset of the striolar hair cell population, we further subclustered striolar hair cells (Figure 3E; Table S2g). This analysis identified a second striolar subcluster marked by *CADM2* (Figure 3F), and a third subcluster with mixed features of striolar and extrastriolar identity, including enrichment of the extrastriolar marker *HTR3B* (Figure 3E), indicating that the striolar compartment is transcriptionally heterogeneous. Module scoring using adult utricle type I and type II gene sets show that the *OCM*^+^ population were more type I-like, while the *CADM2*^+^ population showed a modestly higher type II score, suggesting subtype-associated programs are emerging in the striolar hair cells at W19.

Pseudotime ordering by Monocle3 placed nascent hair cells at the earliest point and striolar hair cells at the latest point along the trajectory, with each transcriptional state occupying a specific pseudotime distribution (Figures 3G-I; Table S4b). Genes that varied dynamically across pseudotime showed a coordinated decrease in early differentiation-associated genes and an increase in genes involved in maturation, function, and stereocilia formation (Figure 3J). Consistent with this progression, Gene Ontology (GO) analysis (Table S5a) of genes upregulated along pseudotime highlighted terms related to cilium development and ion channel activity, supporting progressive functional maturation of hair cells. Conversely, genes downregulated along pseudotime were enriched for processes including axonogenesis and guidance, cell junction assembly, and ribosomal biogenesis (Figure 3K), consistent with patterns described in human neuroepithelial differentiation and suggestive of a shift toward a less biosynthetically active, fate-committed state.^34^ Altogether, we define a developmental trajectory in which fetal utricle hair cells first acquire differentiation-associated programs, followed by spatially patterned striolar and extrastriolar features, while subtype specification is incomplete by W19. These analyses suggest that hair cell differentiation, spatial identity, and subtype maturation unfold along distinct temporal trajectories during human utricle development.

#### Hair cell chromatin accessibility and active transcription factors across development

To investigate whether transcriptional patterns of hair cell maturation are coupled with changes in chromatin landscape, we integrated snRNA-seq and snATAC-seq profiles across the three developmental stages using weighted nearest-neighbour clustering (Figure 4A). This analysis resolved six hair cell clusters spanning nascent, immature, intermediate, mature, extrastriolar, and striolar, broadly consistent with the transcriptomic analysis alone (Table S2h). Differentially accessible regions (DARs) between early developmental populations (nascent, immature, intermediate) and late-stage populations (mature, extrastriolar, striolar) identified stage-enriched peaks; logistic regression (LR) test, *P_adj_* < 0.05 and log_2_FC ≥ 0.5, Table S6a. Motif analysis on these regions revealed early enrichment for basic helix-loop-helix (bHLH) factors associated with neurogenic and sensory differentiation programs (*NEUROD1*, *NEUROG2*, and *ATOH1*), and late enrichment for *POU* family (*POU4F1/2/3*) and homeobox (*LMX1A/B*) motifs (Figure 4B; Table S5b), indicating a shift in chromatin state primed for differentiation toward a fate-restricted regulatory landscape as hair cells mature.

**Figure 4:**
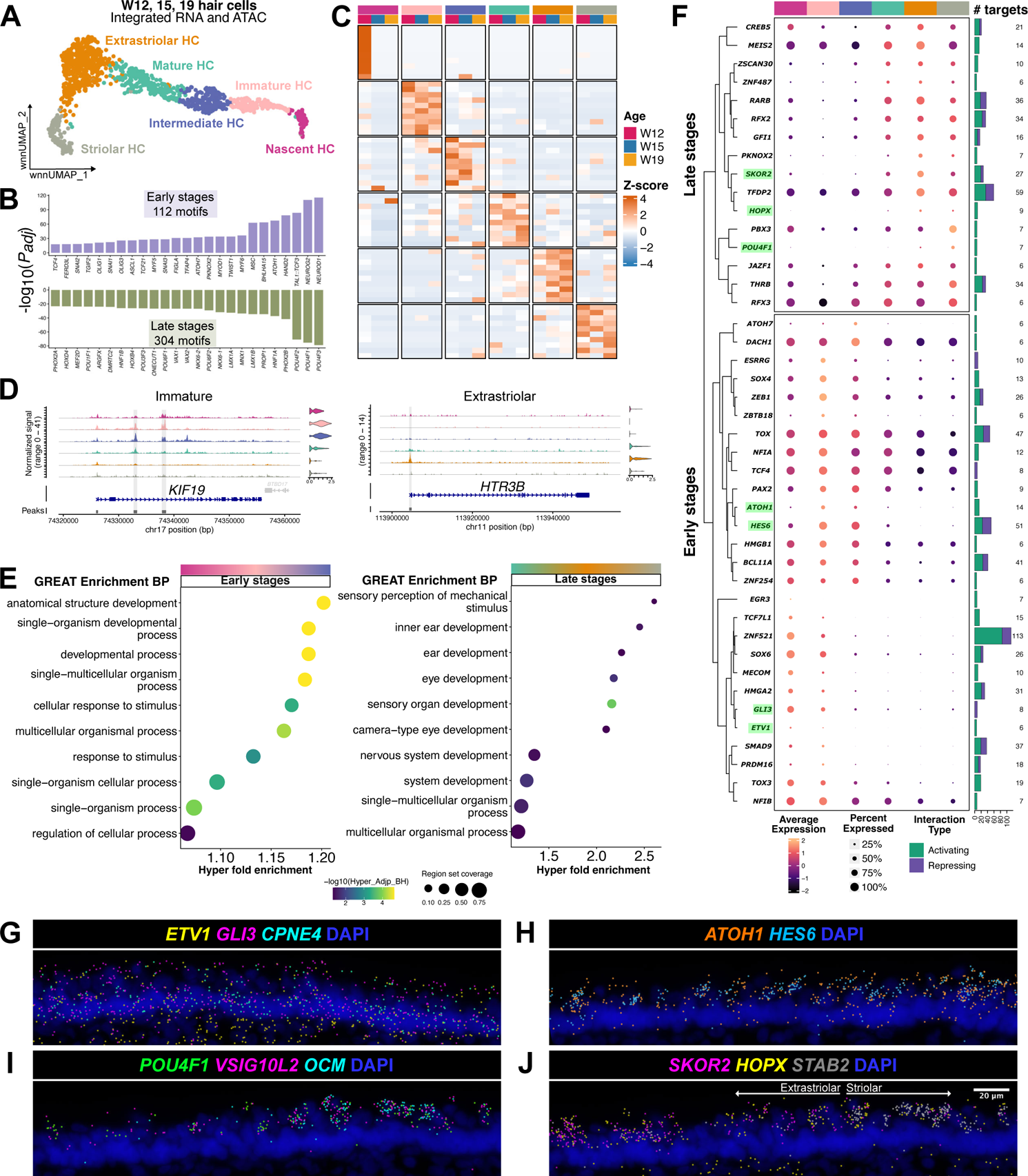
Hair cell chromatin accessibility profiling and active transcription factors. (A) Weighted nearest neighbour (wnn) UMAP projection of integrated snRNA-seq and snATAC-seq data. Cluster annotations are comparable to those obtained from transcriptome-based clustering. (B) Enriched transcription factor motifs at early (nascent, immature, and intermediate clusters) and late (mature, extrastriolar, and striolar clusters) stages of hair cell development. (C) Differentially accessible regions within each cluster and grouped by age. (D) Coverage plots showing accessible peaks around the transcriptional start sites of *KIF19* (immature HCs) and *HTR3B* (extrastriolar HCs). (E) Dot plots showing GREAT-enriched BP terms in early and late stages of hair cell development. (F) Transcription factors with significant inferred downstream regulatory interactions. Plot shows RNA expression (dot colour and size), interaction type (bar colour), and number of target genes (stacked bars). Genes validated in (G-J) are highlighted. (G) Expression of nascent hair cell genes *ETV1* and *GLI3* in supporting cells (*CPNE4^+^*) at W15, n=3. (H) Co-expression of *ATOH1* and *HES6* in immature hair cells, n=3. (I) Co-expression of *POU4F1* with the pan-type I marker *VSIG10L2* and the striolar type I marker *OCM*, n=3. (J) Expression of *SKOR2* and *HOPX* in the extrastriolar hair cells, with *STAB2* marking the striolar region, n=3.

Next, we identified DARs across hair cell populations and developmental stages and linked them to cluster-specific gene expression (Figure 4C; LR test, *P_adj_* < 0.05 and log_2_FC ≥ 0.5, Table S6b). These DARs included an intronic region within *KIF19* that was more accessible in immature hair cells, consistent with a putative enhancer, and increased promoter accessibility at *HTR3B* in extrastriolar hair cells (Figure 4D). To interpret accessibility differences, we used the Genomic Regions Enrichment of Annotations Tool (GREAT) to associate accessible chromatin regions with biological processes (Figure 4E; Table S5c).^35^ Accessible chromatin in early states was primarily associated with cellular development and response to stimulus, whereas late-stage hair cell populations were enriched in inner ear and broader systems development terms. These results indicate that chromatin accessibility patterns encode both developmental progression and emerging spatial subtype identity, consistent with changes in transcriptional programs described earlier.

To identify candidate transcription factors driving hair cell maturation, we inferred a gene regulatory network (GRN) using Pando, which models gene expression based on transcription factor expression with accessibility at their putative binding sites (Figure 4F; Table S7a).^36^ Hierarchical clustering of inferred transcription factor activities separated regulators associated with early hair cell developmental states from those linked to later states (maturation). Among early populations, factors such as *ETV1* and *GLI3* were active in nascent hair cells but were associated with supporting cells by W15 (Figure 4G), suggesting that nascent hair cells retain a progenitor-like regulatory program. Other early-stage regulators included canonical hair cell transcription factors such as *ATOH1* and *HES6* (Figure 4H). In contrast, late-stage transcription factors were associated with functional maturation and survival, including *GFI1* and members of the *RFX* family.

We further identified transcription factors associated with mature hair cell subpopulations. *POU4F1* activity was enriched in type I hair cells and detected in both striolar and extrastriolar regions (Figure 4I). Although POU4F1 is best known for its role in neuronal differentiation and survival of the spiral and vestibular ganglion,^37^ its predicted activity in type I hair cells suggests a potential contribution to the vestibular type I hair cell identity. Additionally, *SKOR2* and *HOPX* emerged as active factors in extrastriolar hair cells (Figure 4J). *Skor2* has been reported in extrastriolar hair cells of zebrafish utricles, and *HOPX* is a Notch-signalling modulator that functions as a transcriptional regulator without direct DNA binding.^38,39^ Collectively, these analyses demonstrate that hair cell maturation is linked to coordinated changes in chromatin accessibility from neurogenic programs towards fate-restricted states, and nominate candidate transcription factors that may contribute to human vestibular hair cell development and the acquisition of subtype and spatial identities.

### Supporting cells exhibit spatial distinction and proliferative decline during development

Supporting cells comprise the largest epithelial population and are key for homeostatic maintenance of the sensory epithelium. Spatial patterning of supporting cells has been described in mice and humans, and we annotated striolar and extrastriolar populations at each age using *TECTB* and *FRZB*, respectively (Figures 5A and C, S4A and B). Differential expression analysis between these regions identified upregulated genes that were age-specific and region-enriched, confirming that striolar and extrastriolar supporting cells maintain distinct transcriptional profiles (Figures 5B and D; *P_adj_* < 0.05, log_2_FC ≥ 1; Table S2i). Across W12, W15, and W19, 166 genes were shared among striolar supporting cells, whereas only 35 were shared among extrastriolar supporting cells (Figure 5E). Striolar supporting cells consistently exhibited a larger set of region-enriched DEGs than extrastriolar supporting cells at all ages, and W19 extrastriolar supporting cells displayed more than twice the number of DEGs observed at W12 or W15, suggesting that the striolar identity is established relatively early, while the extrastriolar supporting cells undergo more dynamic transcriptional changes later in development.

**Figure 5:**
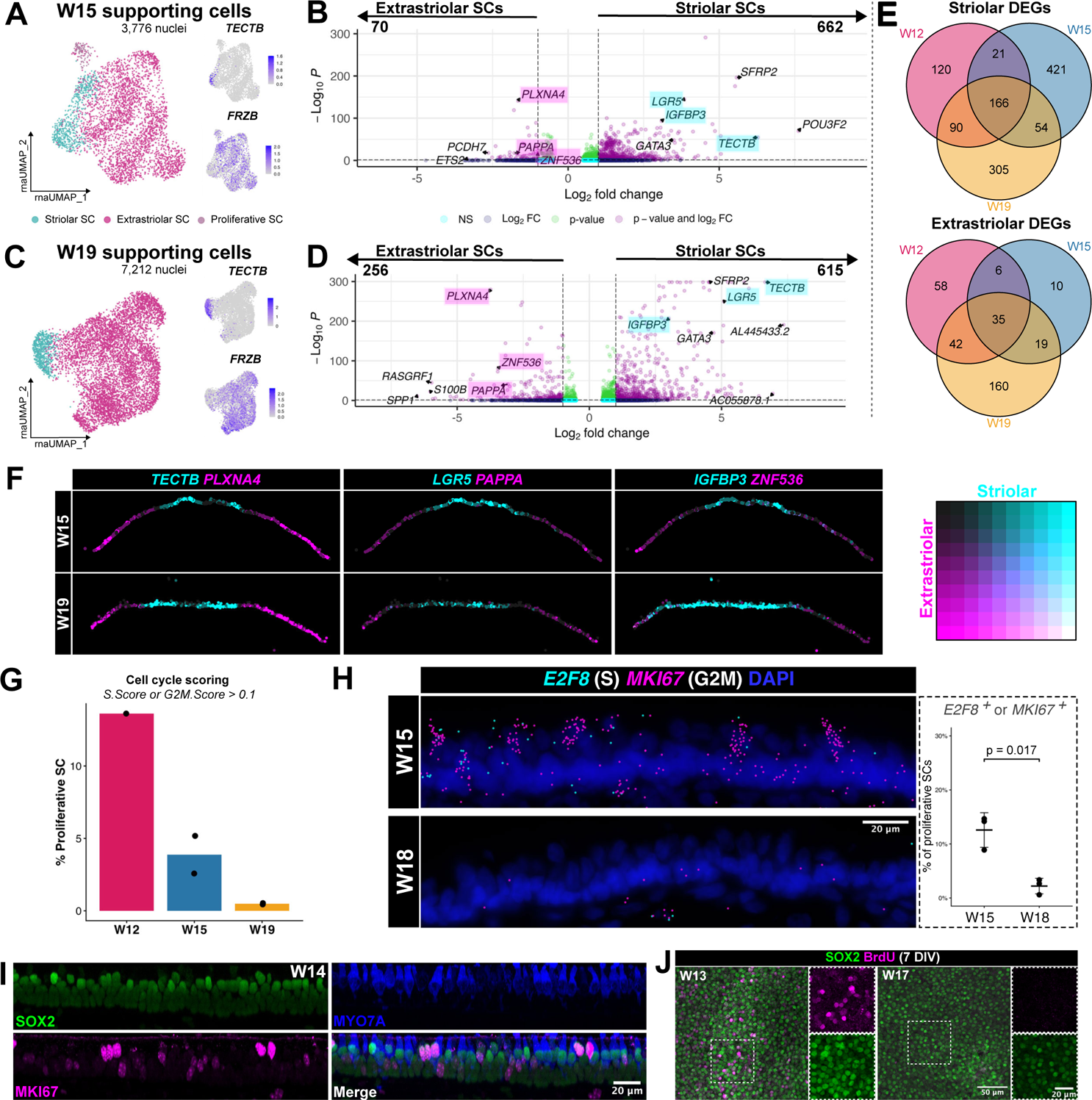
Region-specific transcriptomic differences and proliferation of supporting cells. (A) UMAP projection of W15 striolar and extrastriolar supporting cells (n=2). Insets show expression of striolar (*TECTB*) and extrastriolar (*FRZB*) markers. (B) Volcano plot of differentially expressed genes between striolar and extrastriolar supporting cells at W15. Genes validated in (F) are highlighted. (C) UMAP projection of W19 striolar and extrastriolar supporting cells (n=2). Insets show expression of striolar (*TECTB*) and extrastriolar (*FRZB*) markers. (D) Volcano plot of differentially expressed genes between striolar and extrastriolar supporting cells at W19. Genes validated in (F) are highlighted. (E) Venn diagrams showing the number of DEGs in striolar and extrastriolar supporting cells and their overlap across W12, W15, and W19. (F) Spatial feature plots showing the expression of striolar (*TECTB, LGR5,* and *IGFBP3*) and extrastriolar genes (*PLXNA4, PAPPA,* and *ZNF536*) in W15 and W18 supporting cells, n=3 each. (G) Proportion of proliferating supporting cells at each gestational age derived from cell cycle scoring analysis (S Score or G2M Score > 0.1). Each dot represents one sample. (H) *In situ* validation of *E2F8* and *MKI67* expression at W15 and W18 (n=3), Scale bar: 20 µm. Inset shows quantification of proliferative supporting cells (*E2F8^+^* or *MKI67^+^*) in the Xenium dataset, values represent mean ± standard deviation. (I) Immunohistochemical staining of SOX2 (green), MKI67 (magenta), and MYO7A (blue) in a W14 utricle. Scale bar: 20 µm. (J) BrdU labelling of supporting cells undergoing DNA synthesis after seven days *in vitro* in W13 and W17 utricle explants, n=1 per stage. Scale bar: 50 µm, 20 µm for insets.

Spatial transcriptomics validated regionally enriched supporting cell markers (Figure 5F). *LGR5* and *IGFBP3* were enriched in striolar supporting cells, while *PLXNA4*, *PAPPA*, and *ZNF536* were enriched in the extrastriolar supporting cells. *LGR5* is a Wnt target gene linked to supporting cell-mediated hair cell regeneration, and Plexin A4 (*PLXNA4*) participates in axon guidance via semaphorin-plexin signalling. *ZNF536*, in the extrastriolar supporting cells, encodes a zinc-finger protein that represses retinoic acid-induced transcription, whereas striolar supporting cells express the retinoic acid degradation enzyme *CYP26B1* critical for striolar formation,^40^ consistent with spatially patterned regulation of retinoic acid signalling across the striolar-extrastriolar axis. These data confirm that fetal utricle supporting cells are both transcriptionally and spatially patterned.

We subclustered supporting cells at each age and identified age-specific striolar and extrastriolar populations (Figure S4C). *LGR5* remained enriched in striolar supporting cells, although its expression markedly decreased with increasing gestational age (Figure S4D). Among the extrastriolar supporting cells, four recurrent populations were identified at each age, though they exhibited moderate transcriptional similarity between clusters (Figure S4C; *P_adj_* < 0.05, log_2_FC ≥ 1; Table S2j-l). We validated a subset of supporting cells enriched for *ROBO2* across all ages (Figures S4C and E). *ROBO2* encodes an axon guidance receptor required for proper positioning of primary auditory neuron-hair cell innervations,^41^ suggesting that this subset of *ROBO2*^+^ extrastriolar supporting cells may have a specialized role associated with neurite growth in the utricle sensory epithelium. To map these supporting cell populations *in situ*, we applied Robust Cell Type Decomposition (RCTD) to the Xenium dataset using transcriptomic annotations as a reference.^42^ Although most extrastriolar populations intermingled throughout the extrastriolar region, ExStriolar_SC-3 consistently localized to the sensory epithelial border (Figure S4E). This border-associated population was enriched for transitional epithelial cell markers such as *OC90*,^43^ supporting its identity as a peripheral supporting cell at the edge of the sensory epithelium.

In mice, supporting cells serve as hair cell progenitors and are mitotically active during utricle development, including during the later phases of development.^44^ To examine their proliferative dynamics, we performed cell cycle scoring on supporting cells at each age and observed a marked decrease in mitotically active supporting cells (S or G2/M phases) with increasing gestational age (Figure 5G). Spatial validation using *E2F8* (S phase) and *MKI67* (G2/M phase) confirmed that dividing supporting cells were substantially more abundant at W15 than W18 (Figure 5H; Table S1g). Proliferative supporting cells were not confined to striolar or extrastriolar domains (Figure S4F), but were enriched in the apical region of the sensory epithelium, consistent with interkinetic nuclear migration during epithelial cell cycle progression (Figure 5I).^45^ Explant culture of fetal utricles in BrdU-supplemented media further confirmed active DNA synthesis in supporting cells and revealed a progressive decline in BrdU incorporation with gestational age (Figure 5J). Collectively, these data indicate that supporting cells transition from a proliferative developmental state toward a more quiescent state during the gestational window examined, in parallel with the emergence of spatially distinct striolar and extrastriolar programs.

#### The transitional epithelium displays spatially ordered heterogeneity

Transitional epithelial cells are nonsensory cells located at the periphery of the utricle’s sensory epithelium, forming a boundary between the sensory epithelium and the epithelial roof. To investigate their heterogeneity, we reannotated transitional epithelial cells at W15 and W19 and reintegrated them for analysis, yielding 765 nuclei. We resolved four subpopulations, each marked by cluster-defining genes (Figures 6A and B; *P_adj_* < 0.05, log_2_FC ≥ 1; Table S2m). We validated these markers *in situ* alongside *SOX2*, which is confined to the sensory epithelium, and *CNMD,* which is selective for the transitional epithelium (Figure 6C). Marker expression patterns revealed that each subpopulation occupies a distinct spatial domain, with TEC-1 closest to the sensory epithelium and TEC-4 positioned most peripherally (Figure 6D).

**Figure 6:**
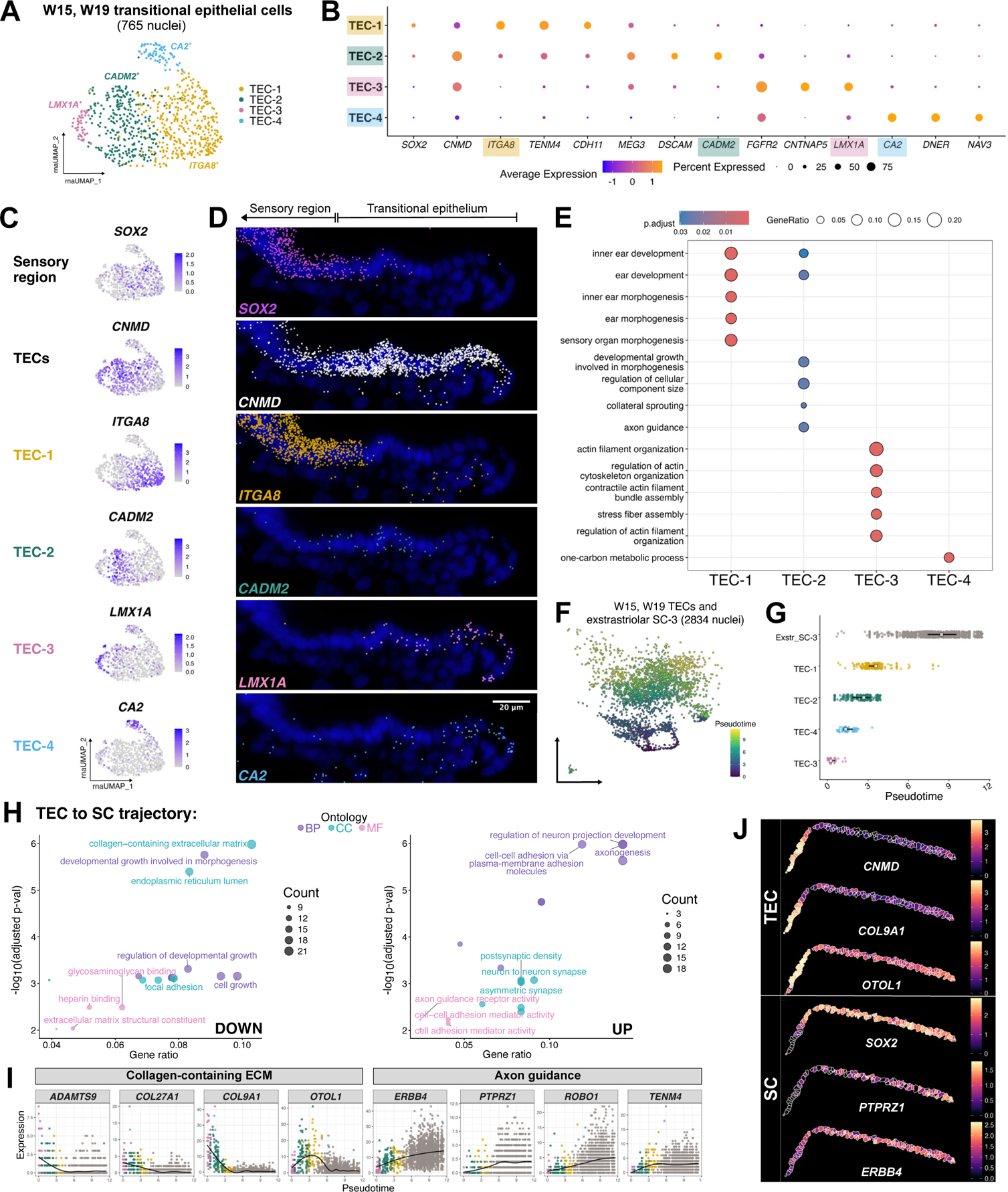
Transitional epithelial cells exhibit spatially distinct gene expression patterns. (A) UMAP projection of integrated W15 and W19 transitional epithelial cells showing four subpopulations, n=4. (B) Dot plot of genes enriched in each cluster. Genes validated in (D) are highlighted. (C) Feature plots of enriched genes in each transitional epithelial cell subpopulation, along with *SOX2* and *CNMD* as markers of the sensory region and transitional epithelium, respectively. (D) *In situ* validation of genes shown in (C). Scale bar: 20 µm. (E) GO BP terms for each subpopulation. (F) UMAP projection of transitional epithelial cells together with Exstr_SC-3 clusters from W15 and W19 supporting cell analysis, coloured by pseudotime. (G) Distribution of pseudotime values by cluster. White squares indicate median pseudotime value; black lines indicate interquartile range. (H) Top three GO terms for BP, CC, and MF enriched among genes upregulated or downregulated along the transitional epithelial cell-to-supporting cell trajectory. (I) Expression of genes inferred to change across pseudotime, grouped into collagen-containing ECM and axon guidance categories. (J) Spatial feature plots of selected genes from (I), n=3.

Cells in TEC-1 showed the greatest transcriptional similarity to supporting cells, including shared *ITGA8* expression and high *SOX2* levels, suggesting a potential intermediate state between transitional and supporting cell identities. TEC-2, marked by *CADM2*, was positioned adjacent to TEC-1. GO analysis indicated that TEC-1 and TEC-2 were the only transitional epithelial cell subpopulations enriched for inner ear development and morphogenesis terms (Figure 6E). Given prior evidence that transitional epithelial cells can act as precursors of supporting cells and contribute to expansion of the sensory epithelium,^46^ these two clusters likely represent the epithelial states with greatest potential to acquire supporting cell features. TEC-3 was marked by *LMX1A*, a transcription factor critical for early inner ear development, and was enriched for actin assembly and organization processes (Figure 6E).^46^ This is consistent with observations in the mouse utricle, where vestibular sensory domains are bounded by regions of *Lmx1a*^+^ transitional epithelial cells and actin cables,^46,47^ suggesting a boundary-forming role for this population. TEC-4, which occupied the most peripheral position, specifically expressed carbonic anhydrases (*CA1/2/3/14*). The enrichment of one-carbon metabolic processes and the known association of carbonic anhydrase activity with vestibular dark cells^48^ point to TEC-4 as a more metabolically specialized, epithelial roof-associated state.

To assess transcriptional continuity with peripheral supporting cells, we integrated transitional epithelial cells with Exstr_SC-3, the most peripheral population, from both ages and performed trajectory inference (Figure 6F; Table S4c). This analysis inferred a trajectory extending from TEC-3 toward Exstr_SC-3 (Figure 6G). GO analysis of genes that changed along pseudotime showed downregulation of genes involved in collagen-containing extracellular matrix (ECM) programs and concomitant enrichment of axon guidance processes (Figures 6H and I; Table S5e). Consistent with this progression, the expression of *COL9A1*, an alpha chain of type IX collagen, and *OTOL1*, an inner-ear specific collagen-like protein, decreased in a gradient from the peripheral transitional epithelium toward the supporting cell compartment. In contrast, gradients in *PTPRZ1* and *ERBB4* expression increased toward supporting cells (Figure 6J). These opposing gradients indicate a coordinated remodelling of functional programs along the transitional epithelium-to-supporting cell continuum. Overall, these findings delineate spatially ordered transitional epithelial states along the sensory epithelium border and support a model of progressive molecular continuity between transitional epithelium cells and peripheral supporting cell identity.

### Distinct epigenomic and signalling programs between epithelial compartments

Having defined the transcriptional and spatial identities of the sensory epithelium, we leveraged paired multiome data to define the regulatory and signalling features that distinguish the cell types of the utricle. Transcriptome-based annotations were transferred onto the multiomic dataset and epigenomic projections to first identify DARs in each cell type (Figures 7A-C; LR test, *P_adj_* < 0.05 and log_2_FC ≥ 0.5, Table S6c). Hair cells, supporting cells, and transitional epithelial cells formed distinct clusters, and W15 and W19 nuclei were partially separated in chromatin accessibility, likely due to age-associated epigenomic differences. Differentially accessible linked peaks between W15 and W19 (LR test, *P_adj_* < 0.05 and log_2_FC ≥ 1, Table S6d) were relatively sparse in hair cells and transitional epithelial cells, indicating comparatively stable chromatin landscapes, whereas supporting cells showed the largest number and effect size of linked peaks, pointing to more extensive chromatin remodelling between W15 and W19 (Figure S5A). This interval notably coincides with a decline in supporting cell proliferation and a reduction in *LGR5* expression, suggesting that the loss of proliferative and regeneration-associated features may be associated with remodelling of the supporting cell chromatin landscape.

**Figure 7:**
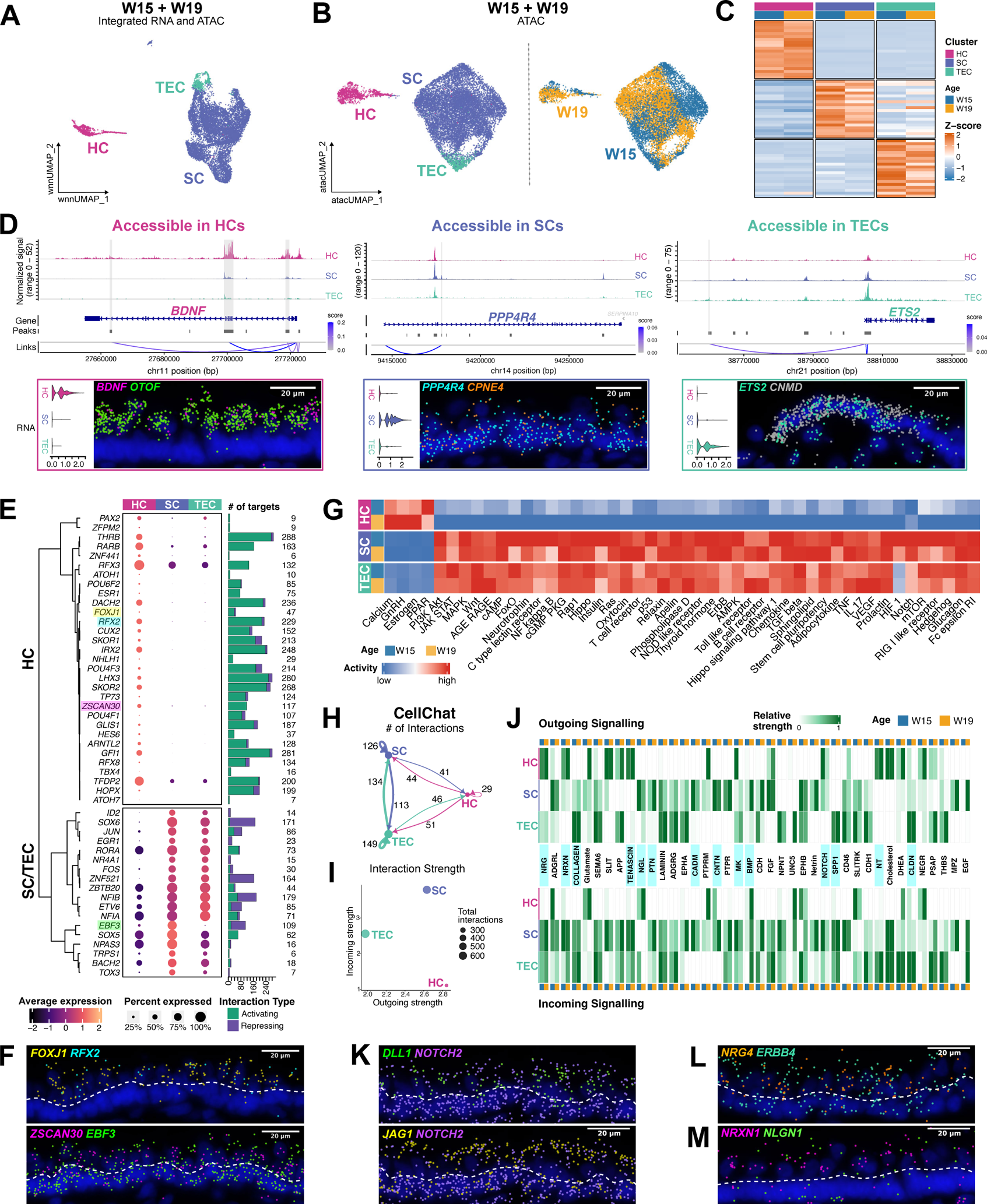
Multiome profiling and intercellular communication between sensory epithelial cell types. (A) wnn UMAP projection of integrated W15 and W19 multiomic data (n=2 per stage). (B) UMAP projection of snATAC-seq data coloured by cluster and stage. (C) Heatmap of DARs within each cell type and grouped by age. (D) Coverage plots of accessible peaks linked to cell type-specific gene expression. Insets depict violin plots of gene expression across cell types and *in situ* validation (W18, n=3) of each gene. Scale bar: 20 µm. (E) Transcription factors with significant inferred downstream regulatory interactions. Genes validated in (F) are highlighted. (F) Expression of active transcription factors in hair cells and supporting cells (W18, n=3). The dashed line separates hair cell and supporting cell nuclei. Scale bar: 20 µm. (G) KEGG signalling pathway activity across cell types and ages. (H) Number of intercellular interactions between each cell type. (I) Strength of incoming and outgoing interactions. (J) Heatmap of the top 40 signalling pathways shared between W15 and W19, in order of outgoing strength. Genes from highlighted pathways are selected for validation. (K) Expression of Notch ligand and receptors in hair cells and supporting cells (W18, n=3). Scale bar: 20 µm. (L and M) Expression of neuregulin signalling LR pair *NRG4-ERBB4* (L) and neurexin LR pair *NRXN1-NLGN1* (M) (W18, n=3). Scale bar: 20 µm.

GREAT enrichment analysis on DARs identified functionally distinct chromatin landscapes across the main cell types (Figure S5B; Table S5f). Hair cell-accessible peaks were strongly enriched for canonical sensory functions including stereocilia-associated and mechanotransduction programs. Supporting cell peaks were associated with epithelial and developmental processes, and transitional epithelial cell peaks were associated with extracellular and matrix-related functions. We also identified cell type-specific peaks linked to representative marker genes, including putative regulatory regions associated with *BDNF* in hair cells, *PPP4R4* in supporting cells, and *ETS2* in transitional epithelial cells (Figure 7D). These findings demonstrate that each epithelial population is characterized by a distinct chromatin-accessibility profile that reflects its cellular identity and function.

To define candidate transcriptional regulators underlying these identities, we inferred GRNs from integrated gene expression and chromatin accessibility (Figure 7E; Table S7b). Hair cells contained the largest set of active transcription factors, most of which were associated with predicted activating interactions. These included *FOXJ1*, implicated in ciliary development, *RFX2*, associated with developing hair cells,^49,50^ as well as *ZSCAN30*, a zinc finger protein previously uncharacterized in hair cells (Figure 7F). Supporting cells and transitional epithelial cells exhibited fewer active transcription factors overall and were dominated by predicted repressive interactions. Among the 18 transcription factors active in the nonsensory populations, only *EBF3* was specific to supporting cells (Figure 7F). These analyses highlight regulatory programs that distinguish hair cells from nonsensory epithelial populations and suggest a partial regulatory similarity between supporting cells and transitional epithelial cells.

We next quantified KEGG signalling pathway activity between cell types using a previously described analysis^51^ by integrating expression of non-ligand pathway genes with GRN-derived activities of pathway-associated transcription factors (Figure 7G; Table S8a). Hair cells displayed a relatively restricted signalling-response profile, with elevated activity in only four pathways: PPAR signalling at W15; and calcium, gonadotropin-releasing hormone (GnRH), and estrogen signalling at W19. In contrast, supporting cells and transitional epithelial cells exhibited broadly similar and more extensive activity, indicating that they are the principal signalling-responsive populations in the fetal utricle. Consistent with these findings, analysis of intercellular communication with CellChat^52^ identified hair cells as having the fewest ligand-receptor (L-R) interactions, the weakest incoming interaction strength via lowest total inferred signalling received, and the strongest outgoing interaction strength via highest total inferred signalling sent (Figures 7H and I). Supporting cells and transitional epithelial cells showed higher numbers of L-R interactions and stronger incoming interaction strength, and supporting cells also displayed strong outgoing signalling.

We identified 63 predicted active signalling pathways among sensory epithelial cell types, 50 of which were shared between W15 and W19 (truncated mean algorithm, Table S8b). From these, we selected representative L-R pairs for spatial transcriptomic validation (Figure 7J, Table S8c). As the Xenium panel captured only a subset of CellChat-related genes, CellChat analysis on the spatial data was used specifically to confirm LR interactions supported by spatial proximity (Figure S5C). Spatial validation confirmed the expression of Notch ligands *DLL1* in hair cells and *JAG1* in supporting cells, together with their receptor *NOTCH2*, corresponding to Notch-mediated communication in the utricle (Figure 7K). Additional inferred programs include pleiotrophin (PTN) signalling within supporting cells (*PTN-PTPRZ1*), involved in cell proliferation and development and corroborates findings from the mouse utricle.^53,54^ Neural development pathways, including neuregulin (NRG) and neurexin (NRXN) signalling, showed a pronounced hair cell-to-supporting cell pattern. We validated *NRG4* ligand in hair cells and its receptor *ERBB4* in supporting cells, as well as *NRXN1* in hair cells and its receptor *NLGN1* in supporting cells (Figures 7L and M). Matrix and adhesion-associated processes, including collagen, tenascin, and cell adhesion molecule (CADM) pathways, were most prominent in supporting cells and transitional epithelial cells. Collectively, these multiomic and pathway analyses reveal distinct chromatin accessibility states and communication programs in the cell types of the utricle sensory epithelium, highlighting supporting cells as a major signalling hub that both receives and relays molecular signals in the sensory epithelium, transitional epithelial cells participating broadly in ECM programs, and hair cells occupying a restricted signal-responsive state.

## DISCUSSION

In this study, we characterized the molecular, spatial, and regulatory programs that shape organ architecture and development of the human utricle sensory epithelium across fetal gestation. By integrating single-nucleus multiome sequencing with imaging-based spatial transcriptomics, we linked cell type-specific transcriptional and epigenomic states with spatially patterned gene expression. We further identified candidate regulatory and signalling programs across epithelial populations, revealing an active remodelling molecular landscape in the developmental window analyzed.

A central finding is that human utricle hair cell development occurs through staged acquisition of differentiation, regional identity, and subtype-specific maturation programs. Rather than maturing uniformly, fetal hair cells progress through transcriptionally distinct states while retaining immature developmental features across the gestational window examined. At W15, differentiation-associated hair cell states were broadly distributed across the sensory epithelium, whereas *OCM* expression already marks an emerging striolar domain, suggesting that spatial identity can be established before hair cell maturation. By W19, *TMC2*^+^ hair cells retained *SOX2* expression and showed continued overlap of canonical adult type I and type II hair cell markers, with only the striolar type I identity clearly resolved. These results support a model in which spatial identity in the human utricle is established before complete transcriptional specification of hair cell subtypes.

The expression of *LGR5* in fetal striolar supporting cells is a notable feature of the developing human utricle, particularly given that *LGR5*^+^ cells have not been reported in available adult human utricle single-cell studies. *Lgr5*^+^ supporting cells in the cochlea and vestibular end organs have been characterized as a *Wnt*-responsive, stem cell-like population capable of regenerating both hair cell subtypes in the neonatal mouse utricle.^55–57^ Its expression across these fetal stages suggests that, similar to mice, *WNT*-responsive supporting cell states are retained in a spatially restricted manner during this period of human utricle development, though it is steadily decreasing with maturation. In parallel, supporting cells showed a progressive decline in proliferation and the most pronounced epigenomic changes among sensory epithelial cell types between W15 and W19, pointing to substantial regulatory remodelling during the same window in which proliferative capacity decreases. Because supporting cell regenerative competence has been linked to chromatin accessibility and proliferation status,^44,58^ these findings indicate that developmental progression across this gestational period is linked to a gradual reduction in regeneration-associated features in the fetal utricle.

Comparisons with murine utricle development further indicate that maturation of the human vestibular sensory epithelium remains ongoing at W19. Prior work^59^ approximates human gestational W12, W15, and W19 to mouse E15, E19, and P6, respectively, providing a broad window for cross-species staging. Within this time frame, we observed progressive molecular changes in hair cells and supporting cells consistent with a conserved maturational trajectory. For example, supporting cell proliferation declined across fetal ages, similar to the reduction of mitotic activity reported in mice during the first postnatal week.^15,44^ However, the incomplete specification of hair cell subtypes and continued *SOX2* expression in *TMC2*^+^ hair cells indicate ongoing maturation. Persistent *LGR5* expression in a subset of striolar supporting cells further suggest a temporal extension of supporting cell maturation, as *Lgr5* was reported to be absent in the striolar supporting cells of the mouse utricle by P5.^55^ Collectively, these observations support a model in which human fetal utricle development follows a broadly conserved maturational sequence relative to mouse, while hair cell subtype specification and supporting cell maturation continue to progress through at least gestational W19.

Our data also demonstrate that transitional epithelial cells are not a transcriptionally homogeneous nonsensory population but instead comprise spatially ordered states along the sensory epithelial border. Transitional epithelial cells subpopulations exhibit either supporting cell-like or roof-like features consistent with their positions along the sensory-nonsensory interface, and trajectory analysis linking transitional epithelial cells with peripheral extrastriolar supporting cells reveals molecular continuity between these populations. This supports a model in which the transitional epithelium is a spatially organized boundary with cell states that bridge the peripheral supporting cell identity to the epithelial roof.

Multiomic analyses further identified changes in the regulatory landscape that accompanies both hair cell maturation and cell type identity. In hair cells, chromatin accessibility transitions from early neurogenic and sensory progenitor motifs towards motifs associated with later differentiation, function, and regional identity, which is consistent with a shift from broad developmental competence to fate-restricted regulatory states. Across the sensory epithelium, hair cells contained the largest set of predicted active transcription factors, mainly associated with activating regulatory interactions, whereas supporting cells and transitional epithelial cells shared overlapping sets of regulators dominated by predicted repressive interactions. These patterns suggest that hair cell maturation involves activation of differentiation programs, while supporting cells and transitional epithelial cells are maintained by regulatory circuitry that stabilizes the nonsensory cell identity.

Within this framework, *POU4F1* emerges as a key transcription factor enriched in type I hair cells. Given that mature utricles spontaneously regenerate type II-like hair cells,^60–63^ the selective association of *POU4F1* with type I hair cells raises the possibility that it contributes to subtype-specific fate stabilization or could act as a barrier to regenerating type I hair cells. In addition, our signalling pathway analyses identify nonsensory populations as major hubs for signalling, with prominent interactions among supporting cells and transitional epithelial cells within the sensory epithelium. Nonsensory cells displayed broad signal-responsiveness and extensive ligand-receptor connectivity, while hair cells exhibited a restricted signal-response profile. Despite having fewer overall interactions, hair cells showed relatively strong outgoing communication strength, suggesting that they act primarily as signal-sending rather than signal-receiving cells within the sensory epithelium. These observations support a view of fetal utricle maturation as a coordinated process in which cell-intrinsic regulatory specialization in hair cells is coupled to extensive intercellular communication orchestrated by nonsensory populations.

In summary, our study fills major gaps in our understanding of spatial, transcriptomic and epigenomic profiles during the development of one of the most understudied human organs, providing a reference for understanding vestibular disorders and uncovering cell states and regulatory pathways that could be targeted in future regenerative strategies.

### Limitations of the study

The interpretation of this study should be guided by some considerations: first, the W12 dataset was derived from a single fetal sample and was therefore used to contextualize development rather than to support age-specific statistical comparisons. Second, although the targeted Xenium panel provided high-resolution spatial validation of selected genes, it is not an unbiased reference of the whole transcriptome. Therefore, RCTD label transfer and CellChat inference from the Xenium data should be interpreted as targeted spatial support. Both considerations clarify the scope within which the findings should be interpreted and do not affect the utility or potential of our datasets.

## RESOURCE AVAILABILITY

### Lead contact

Further information and requests for resources and reagents should be directed to and will be fulfilled by the lead contact, Alain Dabdoub.

### Materials availability

This study did not generate any unique reagents. Researchers interested in accessing future collected samples can contact the corresponding authors.

### Data and code availability

- Raw sequencing data (10x Multiome snRNA-seq/snATAC-seq) and Xenium spatial transcriptomic outputs have been deposited in the NCBI Gene Expression Omnibus database under accession code GSE######. Accession numbers for all raw datasets are also listed in the key resources table. Processed data have been deposited in the gEAR portal (umgear.org).
- This study does not report original code.
- Any additional information required to reanalyze the data in this paper is available from the corresponding authors.

## ACKNOWLEDGEMENTS

We thank the members of the Dabdoub lab, particularly Boaz Ehiogu, for sharing and providing guidance with the gene regulatory modules inference pipeline, and Dr. Matsya R. Thulasiram for helpful discussion and support. We also thank the donors, RCWIH BioBank, the Lunenfeld-Tanenbaum Research Institute, and the Mount Sinai Hospital/University Health Network (UHN) Department of Obstetrics and Gynaecology for the human specimens used in this study (biobank.lunenfeld.ca). We thank the Princess Margaret Genomics Centre, and the UHN Bioinformatics and High-Performance Computing Core and Pathology Research Program Laboratory for providing high-throughput sequencing, bioinformatic services, and spatial transcriptomics assistance. We are grateful for the Ottawa Bioinformatics Core Facility, especially Christopher Porter and Asma Bankapur, for bioinformatics support. We would also like to thank the Harvard Chan Bioinformatics Core, Harvard T.H. Chan School of Public Health, Boston, MA, RRID:SCR_025373, for feedback on the snATAC-seq pipeline. We also acknowledge Compute Ontario (computeontario.ca) and the Digital Research Alliance of Canada (alliancecan.ca) for providing hardware, storage, and computing resources, as well as the imaging facilities at Sunnybrook Research Institute and SickKids Research Institute. We extend our thanks to the gEAR team, particularly Daniel Lesperance and Ricky S. Adkins, for their support in uploading datasets onto the platform, and we thank Drs. Jennifer Stone and Yong Zeng for their comments on the manuscript. This work was supported by funding from the Canadian Institutes of Health Research – CIHR PJT-183579 (E.L. and A.D.); Hearing Health Foundation’s Hearing Restoration Project (A.D.); Canadian Graduate Scholarship – Doctoral Program from CIHR (W.L.); Canadian Graduate Scholarship – Master’s Program from CIHR (W.L.); Raymond H.W. Ng Graduate and Doctoral Awards, as well as Harry Barberian Research Award from the Department of Otolaryngology at the University of Toronto (W.L.); Ontario Graduate Scholarship – Doctoral Award from the Government of Ontario (W.L.); Japan Society for the Promotion of Science – JSPS Postdoctoral Fellowship (R.Y.); and the Michael and Sonja Koerner Charitable Foundation (A.D.).

## AUTHOR CONTRIBUTIONS

E.L. and A.D. conceived the study; W.L. and E.L. collected the samples; E.L. generated the multiomic datasets, and W.L., R.Y., and E.L. generated the spatial transcriptomic datasets; W.L. performed the computational analyses with support from R.Y. and E.L; W.L. performed sample preparation for spatial transcriptomics, explant cultures, immunohistochemistry, and imaging; W.L., R.Y., E.L., and A.D. analyzed and interpreted the data; W.L. wrote the original draft; all authors revised and edited the manuscript; E.L. and A.D. supervised the study. All authors read and approved the manuscript.

## DECLARATION OF GENERATIVE AI AND AI-ASSISTED TECHNOLOGIES

During the preparation of this work, the authors used ChatGPT (v.5.5) to partially assist in developing visualization approaches for snRNA-seq and snATAC-seq data. The authors reviewed and edited all AI-assisted outputs and take full responsibility for the content of the publication.

## SUPPLEMENTARY FILE INFORMATION

**Supplementary Table S1: Sample metadata, sequencing metrics, disease-associated gene expression, and proliferating supporting cell quantification**

Table provides details of (a) all samples used in this study for snMultiome, Xenium, tissue experiments and imaging. Sample information includes age, sex, snMultiome (GEX and ATAC) library construction, sequencing details, and quality control metrics, Xenium sample preparation and output, and experiment details, related to all figures; (b) percentage of utricle epithelium cells that express genes associated with vestibular impairment, related to Figure S1A, (c) syndromic and non-syndromic hearing loss related to Figure S1B; (d) correlation of hair cell differentiation states with spatial identity in W15 Xenium hair cells and statistical summary, related to Figure 2H and I. (e) quantification of *VSIG10L2* and *CALB1* expression in W18 hair cells, related to Figure S2E; (f) adult utricle hair cell subtype markers for module score analysis, related to Figures S2F and S3J; and (g) quantification and statistical details for supporting cell proliferation, related to Figure 5H.

**Supplementary Table S2: Differential gene expression analyses**

Table contains all differentially expressed genes (DEGs) for (a) hair cells, supporting cells, and transitional epithelial cells in the integrated W15 and W19 dataset, related to Figures 1F-H; W15 (b), W19 (c), and W12 (d) hair cell clusters, related to Figures 2D-G, L-P, S2B, and S3F; integrated W12, W15, and W19 hair cell clusters (e) prior to and (f) after removal of a high mitochondrial/ribosomal gene expressing cluster, related to Figures S3H and 3D; (g) striolar hair cell subclusters, related to Figure 3E; (h) integrated hair cell clusters following weighted-nearest-neighbour (wnn) clustering, related to Figure 4A; (i) striolar and extrastriolar supporting cells across three gestational ages, related to Figures 5B, D-F, S4B; (j) W15, (k) W19, and (l) W12 supporting cell clusters, related to Figure S4C; and (m) integrated W15 and W19 transitional epithelial cell clusters, related to Figures 6B-D. Enriched genes are determined using Wilcoxon’s rank-sum test with *P_adj_* < 0.05.

**Supplementary Table S3: Custom Xenium gene panel details and cell type annotations**

Table contains a list of genes included in the customized panel (ID V2D67G) for Xenium *In Situ* spatial transcriptomics. Information includes gene symbol, Ensembl ID, probe sets, and cell type annotations. Related to all figures.

**Supplementary Table S4: Pseudotime trajectory analyses using Monocle 3**

Table contains all genes associated with pseudotime trajectories reconstructed using Monocle 3 for (a) W15 hair cells, related to Figures 2C and S2A; (b) integrated W12, W15, and W19 fetal hair cell development, related to Figures 3G-K; and (c) integrated W15 and W19 transitional epithelial cell-to-supporting cell trajectory, related to Figures 6F-J. Directionality of gene expression (up or down) across pseudotime is determined using Spearman’s correlation and included where applicable.

**Supplementary Table S5: Gene ontology (GO), transcription factor motif, and Functional Enrichment of Genomic Regions (GREAT) enrichment analyses**

Table contains (a) enriched GO terms among genes regulated along the fetal hair cell development trajectory, related to Figure 3K; (b) transcription factor motifs enriched in early and late hair cell differentiation stages, related to Figure 4B; (c) GREAT biological process terms for accessible chromatin in early and late hair cell differentiation states, related to Figure 4E; (d) enriched biological process GO terms in the four transitional epithelial cell populations, related to Figure 6E; (e) enriched GO terms for genes regulated along the transitional epithelial cell to supporting cell trajectory, related to Figures 6H and I; and (f) GREAT enrichment results for accessible chromatin in hair cells, supporting cells, and transitional epithelial cells, related to Figure S5B. All analyses were performed using a hypergeometric test against a matched background gene/peak set for GO and motif/GREAT analyses, respectively and adjusted for multiple comparisons using a Benjamini-Hochberg test. Terms with *P_adj_* < 0.05 are reported.

**Supplementary Table S6: Differential accessibility and peak-gene link analyses between hair cell differentiation stages, epithelial cell types, and gestational ages**

Table contains differentially accessible (DA) regions for (a) early *vs* late hair cell differentiation stages, related to Figures 4B and E; (b) integrated W12, W15, and W19 hair cell clusters following wnn clustering, related to Figures 4C and D; (c) integrated W15 and W19 epithelial cell types, related to Figures 7C, D, and S5B; and (d) W15 *vs* W19 comparisons for DA regions between each epithelial cell type, related to Figure S5A. Linked genes are included where applicable. DA regions are determined using a logistic regression test with *P_adj_* < 0.05.

**Supplementary Table S7: Active transcription factors and downstream target genes inferred using Pando gene regulatory network analysis**

Table contains gene regulatory network inferences using Pando, and includes active transcription factors, downstream targets, and predicted regulatory direction for (a) integrated W12, W15, and W19 hair cell subpopulations, related to Figure 4F; and (b) integrated W15 and W19 hair cells, supporting cells, and transitional epithelial cells, related to Figure 7E.

**Supplementary Table S8: KEGG pathway activity and CellChat output**

Table includes (a) KEGG pathway activity quantification across cell types, related to Figure 7G; (b) inferred intercellular communication pathways between cell types based on integrated W15 and W19 transcriptomic data, related to Figure 7J; and (c) significant CellChat-inferred pathways in W15 and W18 Xenium samples, related to Figure S5C.

## METHODS

## EXPERIMENTAL MODEL AND STUDY PARTICIPANT DETAILS

### Ethics approval

De-identified human fetal samples were obtained from the Research Centre for Women’s and Infants’ Health biobank under approval from the Research Ethics Board of Mount Sinai Hospital (ID 20-0003-E) and Sunnybrook Health Sciences Centre (ID 1514). Tissues used in this study are listed in Table S1a.

### Sample collection

Samples were obtained from elective terminations following written informed consent. No compensation was provided. Maternal exclusion criteria included self-reported medical disorders, viral or bacterial infections, and substance use. Fetal exclusion criteria included structural abnormalities, abnormal growth (small or large for gestational age), and exposure to chemical substances. Fetal skulls at target gestational ages were transported on ice in Hanks’ Balanced Salt Solution (HBSS^+/+^; Wisent; #311-512-CL) supplemented with 10mM HEPES (Wisent; #330-050-EL) with a collection-to-lab interval of 1-4 hours. Gestational age was determined based on clinical ultrasound (ACUSON Juniper^TM^ Ultrasound System; Siemens Healthineers) and by using a standard fetal growth table.^64^ Sex was determined by genotyping *ZFX* and *ZFY* from additional tissue from the same specimen.

## METHOD DETAILS

### Bony labyrinth and utricle dissection

Bony labyrinths were dissected from the skull in a black sylgard-coated dish containing ice-cold HBSS^+/+^ with 10mM HEPES. Cartilage overlaying the vestibular region was trimmed with Dumont #3 forceps (Fine Science Tools, 11231-30) to expose the vestibular organs. The vestibular nerve was cut using Dumont #5 forceps (Fine Science Tools, 11251-10), and the sub-epithelial mesenchyme of the utricle was grasped to gently remove the organ from the bony labyrinth. The epithelial roof was carefully removed with Dumont #55 forceps (Fine Science Tools, 11255-20), and otoconia were flushed away using a 1 mL insulin syringe/30 G ½” needle (BD, 324704).

### Utricle explant cultures for *in vitro* proliferation

Utricles were cultured free-floating for 7 days *in vitro* (DIV) in 35 mm dishes (Falcon, 351008) containing 3 mL of DMEM/F-12 GlutaMAX (ThermoFisher Scientific, 10565042) supplemented with 10% fetal bovine serum (Invitrogen, 12485028), 2.5 µg/mL Amphotericin B/fungizone (ThermoFisher Scientific, 15290018), 10 µg/mL ciprofloxacin (Sigma-Aldrich, 17850-5G-F), and 0.35 ng/mL 5-bromo-2’-deoxyuridine (BrdU; BD Biosciences, 550891). Cultures were maintained at 37 °C in a humidified incubator with 5% CO_2_ with complete media changes every other day.

### Histology

#### Fixation and cryosectioning

Utricles were fixed overnight at 4 °C in 4% paraformaldehyde (PFA; Electron Microscopy Sciences, 15710) in phosphate-buffered saline (PBS; Wisent Inc., 311-010-CL). After fixation, samples were washed with 1x PBS at room temperature (RT) and stored at 4 °C until processing. For cryosectioning, tissues were washed in 1x PBS, equilibrated through a sucrose gradient (10%, 20%, 30%; Sigma-Aldrich, S0389), and embedded in Optimal Cutting Temperature compound (OCT; Fisher Healthcare, 23730571) in cryomolds (Tissue-Tek, 4565). Blocks were stored at -80 °C in sealed plastic bags. Sections (10 µm) were cut onto Superfrost Plus Microscope slides (Fisher Scientific, 12-550-15) and stored at -80 °C in sealed slide boxes with desiccant (VWR, 61161-319).

#### Immunofluorescence

Utricle sections or whole mounts were permeabilized for 30 min in 0.5% Triton X-100 (ThermoFisher Scientific, BP151-500) in 1x PBS, then quenched for 30 min in 0.3 M glycine (ThermoFisher Scientific, BP381-1) in permeabilization buffer. For MKI67 and BrdU staining, antigen retrieval was performed with 1 N HCl (diluted from 12 N in distilled water; Fisher Scientific, A144-500) for 30 min at RT, followed by three 15 min washes in distilled water (Invitrogen, 10977-015). Non-specific binding was blocked for 1 h at RT in permeabilization buffer containing 10% Donkey Serum (Sigma-Aldrich, D9663). Primary antibodies were diluted in blocking solution and incubated overnight at 4 °C. The next day, samples were washed for 30 min with permeabilization buffer and incubated for 1 h at RT with Alexa Fluor-conjugated secondary antibodies and DAPI (1 µg/mL, Sigma-Aldrich, D9542) in blocking solution. Samples were then washed for 1 h with 1x PBS and mounted using Fluoromount G (Southern Biotech, 0100-01) or ProLong^TM^ Gold Antifade Mountant (Fisher Scientific, P36930).

#### Antibodies

Primary antibodies used were: mouse anti-MYO7A (1:100, Santa Cruz Biotechnology, sc-74516), rabbit anti-MYO7A (1:1000, Proteus BioScience, 25-6790), rabbit anti-MKI67 (1:250, Abcam, ab15580), rat anti-SOX2 (1:250, eBioscience, 14-9811-80), and mouse anti-BrdU (1:250, BD Pharmingen, 555627). Alexa Fluor-conjugated secondary antibodies used (1:1000; Invitrogen) were: donkey anti-mouse AF488 (A-21202), donkey anti-mouse AF555 (A-31570), donkey anti-rabbit AF555 (A-31572), donkey anti-rabbit AF647 (A-31573), and donkey anti-rat AF488 (A-21208). For hair bundle labelling after secondary antibody incubation, tissues were incubated with Alexa Fluor Plus 647 Phalloidin (1:1000, ThermoFisher Scientific, A30107) for 10 min at RT.

#### Imaging

Images were acquired with a Leica DIM6000 SP8 Lightning Confocal STED or STELLARIS 8 FALCON w/STED confocal microscopes using 20x/0.8 NA, 40x/1.3 oil-immersion, and 63x/1.4 oil-immersion objectives. Z-stacks spanning the full depth were collected with Leica LAS X-optimized step sizes of 0.3 to 0.6 µm and rendered as maximum-intensity projections. Images were processed using Fiji (v2.16.0).

### 10x Single-nucleus Multiomics sequencing (snRNA-seq and snATAC-seq)

#### Sensory epithelium dissection

Six utricles (W12, n=1; W15, W19, n=2 each) were dissected and incubated individually for 10 min at 37 °C in prewarmed 2 mg/mL thermolysin from *Geobacillus stearothermophilus* (Sigma-Aldrich, P1512) in HBSS^+/+^ with 10 mM HEPES. Each utricle was incubated in 2 mL enzymatic solution in a 1-dram glass vial with a rubber-lined cap (15x45 mm; Fisher, 03-339-25B). Samples were then transferred to cold HBSS^+/+^ with 10 mM HEPES and 10% FBS to quench enzymatic activity.

The sensory epithelium was gently delaminated from the underlying mesenchyme using a 1 mL insulin syringe/30 G ½’’ needle and transferred to a 1.5 mL DNA LoBind tube (Fisher Scientific, 13-698-791) containing cold HBSS^+/+^ with 10mM HEPES. For W12, left and right sensory epithelia were pooled. Tubes were centrifuged at 300 × *g* for 6 min at 4 °C, supernatant was removed using a 1 mL insulin syringe/30 G ½’’ needle, and pellets were flash frozen and stored in liquid nitrogen until nuclei isolation.

#### Tissue dissociation, single-nucleus isolation, library preparation and sequencing

The 10x Genomics nuclei isolation protocol (Chromium Nuclei Isolation Kit, User Guide CG000505) was modified as previously described by Deng *et al*.^65^ Single-nucleus Multiome (snMultiome) libraries were generated using the 10x Chromium Single Cell Multiome ATAC + Gene Expression (GEX) Kit (10x Genomics, PN 1000282) according to the manufacturer’s instructions. Briefly, nuclei from sensory epithelia were isolated using lysis buffer and gentle trituration with a P1000 pipette for 6 min. Nuclei suspensions were filtered through a 40 µm Flowmi Cell Strainer (Sigma-Aldrich, BAH136800040) and pelleted at 500 × *g* for 3 min. With the exception of W12, pellets were resuspended in debris removal buffer and centrifuged at 700 × *g* for 10 min, then resuspended in a 1:1 mixture of wash buffer and resuspension buffer and centrifuged at 500 × *g* for 10 min. Final pellets were resuspended in resuspension buffer. Approximately 6,000 to 10,000 nuclei were captured per sample with >90% intact nuclei by acridine orange/propidium iodide (AOPI) staining. Library construction followed the Chromium Single Cell Multiome ATAC + GEX User Guide (CG000338). Nuclei suspensions were incubated in transposition mix for 1 h in a C1000 Touch thermal cycler (Bio-Rad) to fragment accessible chromatin and add adapters. Nuclei were loaded onto a 10x Multiome J chip with gel beads and oil to generate gel bead-in-emulsion (GEM), each encapsulating a single nucleus. GEMs were transferred to pre-chilled PCR strip tubes (MJS BioLynx, #US14024700) for first-strand cDNA synthesis and barcoding of mRNA and transposed DNA.

Reactions were quenched by adding Quenching Agent (10x Genomics, 2000269), and samples were stored at -80 °C for <4 weeks before library preparation.

Pre-amplification products were recovered with Recovery Agent (10x Genomics, PN 220016), purified with a Silane DynaBead mix, and split to generate separate gene expression (GEX) and ATAC libraries. For GEX libraries, purified material was amplified for 7 cycles and cleaned with SPRIselect beads (Beckman Coulter, cat. no. B23318). For ATAC libraries, material was amplified for 9 cycles and similarly cleaned with SPRIselect beads. Library size and quality were assessed on a Bioanalyzer (Agilent Technologies). During sample-indexing PCR, GEX and ATAC libraries were amplified for 12 and 9 cycles, respectively, followed by a final SPRIselect beads clean-up. GEX and ATAC libraries were sequenced separately on Illumina’s NovaSeq X (W12 sample 1, W15 sample 1) or NovaSeq 6000 (W15 sample 2; W19 sample 1 and 2).

### 10x Genomics Xenium spatial transcriptomics

#### Sample preparation and cryosectioning

All instruments and surfaces were cleaned with RNase Away Surface Decontaminant (Thermo Scientific, 7002PK) before tissue processing. W15 and W18 utricles (n=3 each) were dissected and immediately embedded in OCT in a cryomold, oriented in the medial-lateral plane. Blocks were frozen according to the Xenium Fresh Frozen Tissue Preparation Handbook (10x Genomics, CG000579) and stored at -80 °C for up to 15 months. Tissue was cryosectioned at 10 µm, collected onto Xenium slides (10x Genomics, PN3000941), and stored at -80 °C for up to three days before processing.

#### Custom panel preparation and tissue processing for Xenium In Situ Spatial Transcriptomics

A custom gene panel was designed based on the single-nucleus multiome analysis and included known and newly identified cell type- and region-specific markers, transcription factors from gene regulatory network analysis, and ligand-receptor pairs inferred by CellChat. A lyophilized four-reaction panel (Design ID V2D67G) was generated and stored at -20 °C.

Tissue sections were processed according to Xenium In Situ Demonstrated Protocols (10x Genomics, CG000581; CG000582) using manufacturer-provided buffers. Briefly, Xenium slides were fixed and permeabilized at RT, then hybridized overnight at 50 °C with the custom panel, followed by post-hybridization washes (37 °C for 30 min). Sections then underwent ligation (37 °C for 2 h) and amplification (30 °C for 2 h). Cell segmentation staining was then performed according to the protocol (10x Genomics, CG000749), by incubating sections overnight at 4 °C with the Xenium Multi-Tissue Stain Mix (10x Genomics, PN 2000991) followed by autofluorescence quenching (10 min at RT) and nuclear staining. Slides were then loaded onto Xenium Analyzer (10x Genomics, XETG00082) for transcript decoding and imaging (software v3.2.1.2, analysis v3.2.0.7). After imaging, slides were treated with Quencher Removal Solution and stained with hematoxylin & eosin (H&E) as per the protocol (10x Genomics, CG000613).

## QUANTIFICATION AND STATISTICAL ANALYSIS

### snMultiome data preprocessing and analysis

Sequencing reads for RNA and ATAC were aligned to the human GRCh38 reference using Cell Ranger ARC (v2.0.0-2.0.2). W15 libraries were aggregated with cellranger aggr, downsampling the NovaSeq X run (sample 1) to match the depth of the NovaSeq 6000 run (sample 2). Ambient RNA was removed from the RNA modality using SoupX^66^ (v1.6.2). Downstream transcriptomic analyses were performed in Seurat^20^ (v4.3.0-5.3.0). Doublets were identified with scDblFinder (v1.18.0) and excluded, and nuclei were filtered using quality control parameters listed in Table S1a.

Gene expression data were normalized using SCTransform^67^ (vst method, 3000 variable features). Integration anchors were identified using RPCA followed by SCTransform renormalization and PCA (top 30 PCs). UMAPs were generated from the PCA space. Leiden clustering, with resolutions guided by Clustree^68^ (v0.5.1) up to 0.5, was used to define clusters. Differentially expressed genes were identified using Seurat’s FindAllMarkers (Wilcoxon rank-sum test, *P_adj_* < 0.05), and clusters were annotated by canonical marker expression. For W12, only epithelial cells (*EPCAM^+^*) were retained; mesenchyme (*PDGFRA^+^*), glia (*CDH19^+^*), and pericytes (*RGS5^+^*) were excluded. Seurat’s CellCycleScoring function was used to categorize proliferative cells in the transcriptomic data.

Pseudo-bulk peak calling for snATAC-seq was performed with MACS2^69^ (v2.2.9.1) using transcriptomic-defined clusters. Peaks were normalized and scaled in Signac^70^ (v1.9.0-1.14.0) using term frequency-inverse document frequency (TF-IDF) transformation, and dimension reduction was performed using latent semantic indexing (LSI; 2 to 20 dimensions). Reciprocal LSI was used to define integration anchors for accessibility data, and RNA and ATAC modalities were integrated with weighted-nearest-neighbour (wnn) analysis.

Peak-gene links were inferred with Signac’s LinkPeaks function with default parameters: Pearson correlation method, distance threshold = 500,000 bp, minimum cells = 10, *P* ≤ 0.05. Differentially accessible peaks were identified by logistic regression (LR), testing peaks present in at least 5% of nuclei in either group and defined as significant at log_2_FC ≥ 0.5 and *P_adj_* < 0.05.

### Xenium data preprocessing and analysis

Transcript expression was visualized using Xenium Explorer (10x Genomics, v3.2.0). Transcripts with a Xenium quality score < 20 and empty cells were excluded from downstream analysis. Processed Xenium outputs were then analyzed using the same Seurat workflow as the snRNA-seq in this study. Cell boundaries were defined using the 10x Genomics multimodal cell segmentation algorithm based on multi-channel stain images of cell membranes and intracellular RNA/proteins. Data were normalized and scaled with SCTransform followed by PCA (30 PCs) and UMAP dimensionality reduction. Clusters were identified using Leiden clustering and annotated based on canonical marker expression.

Spatial deconvolution of supporting cell subpopulations was performed using Robust Cell Type Decomposition^42^ (RCTD) on doublet mode suited for the high resolution of Xenium In Situ data. Supporting cells from each Xenium sample were queried using snRNA-seq supporting cell cluster identities as the reference.

Proliferative cells were defined as those expressing at least one transcript of *E2F8* or *MKI67*, and statistical significance between groups was assessed using a two-sided Welch’s t-test. Transcript counts are reported in Table S1f.

### Hair cell subtype module score analysis

To quantify spatial bias in W15 hair cell maturation programs, hair cell maturation-state module scores were calculated in W15 Xenium using Seurat’s AddModuleScore function. Gene sets for each category were derived from immature, intermediate, and mature hair cell DEGs. To assess whether these programs were spatially associated, *OCM* expression was used as a proxy for striolar hair cells, and Spearman correlations were calculated between cell-level *OCM* expression and each maturation state module score within each Xenium sample. Sample-level correlation coefficients were tested against zero, and *P*-values were adjusted across module scores using the Benjamini-Hochberg method.

Hair cell subtype module scores were calculated to compare Type I- and Type II-associated programs. Gene sets were generated using only the top 20 genes for each subtype, ranked by log_2_FC, from Wang *et al.,* 2024 and Luca *et al.,* 2026.^18,71^ The full list of hair cell subtype genes is reported in Table S1e.

### Pseudotime and cell-state ordering using Monocle 3

Pseudotime and trajectory inference were performed using Monocle 3^23^ (v3.1.4.26). Analyses were run separately for hair cells across development and for the transitional epithelial cell-to-supporting cell compartment, following the standard Monocle 3 workflow for dimension reduction, principal graph learning, and pseudotime ordering. Root nodes were selected based on expected lineage origin: nascent/immature hair cells for the hair cell trajectory and transitional epithelial cells for the transitional epithelial cell-to-supporting cell trajectory.

Trajectory-associated genes were identified using Moran’s I statistic on the principal graph, and genes with q < 0.01 were considered significant. Significant genes were classified as increasing (ρ > 0.2) or decreasing (ρ < -0.2) along pseudotime based on Spearman’s rank correlation between normalized gene expression and pseudotime.

### Gene ontology and functional enrichment on genomic regions analyses

GO enrichment analysis was performed on Monocle 3-derived trajectory-associated genes and on genes defining transitional epithelial cell subpopulations using clusterProfiler^72^ (v4.12.6) with human gene annotations from org.Hs.eg.db (v3.19.1). Enriched GO terms were considered significant at Benjamini-Hochberg *P_adj_* < 0.05.

Functional enrichment of accessible chromatin regions was assessed with the Genomic Regions Enrichment of Annotations Tool^35^ (GREAT) via the package rGREAT^73^ (v2.6.0) Differentially accessible peaks for specified cell identities (early *vs* late-stage hair cells, or epithelial cell populations) were used as the foreground, and all other called peaks in the corresponding dataset served as the background. GREAT analysis was run with default parameters, and enriched terms were retained using HyperAdjustedP < 0.05, HyperFoldEnrichment > 1.0, and included at least three foreground gene hits.

### Motif enrichment analysis of hair cell peaks

Motif enrichment analysis was performed using Signac. Transcription factor (TF) position frequency matrices were obtained from the JASPAR2020 database (v.0.99.10) with TFBSTools (v1.42.0), restricted to *Homo sapiens* motifs. Differentially accessible peaks between early- and late-stage hair cell clusters were identified using the LR method (log_2_FC ≥ |0.25| and *P_adj_* < 0.05) and used as input to FindMotifs for motif enrichment.

### Gene regulatory network inference

Gene regulatory networks (GRNs) for hair cells and sensory epithelial cell types were inferred using Pando^36^ (v1.1.1). Pando objects were initialized from SCT-normalized gene expression and called peaks. TF motif occurrence was annotated on the hg38 genome using position frequency matrices and TF-motif mappings provided with Pando.

Network inference was restricted to genes differentially expressed among hair cell states or epithelial cell types. Peak-to-gene associations were computed using the GREAT-based linking strategy in Pando, and TF-target associations were retained using a TF correlation threshold of 0.2. Regulatory modules were identified using Pando’s default thresholds: *P_adj_* ≤ 0.05, R^2^ ≥ 0.1, minimum number of variables = 10, minimum genes per module = 5). Positive and negative target gene modules were extracted for each TF.

### KEGG pathway activity quantification and CellChat intercellular communication analysis

KEGG pathway activity scoring was adapted from Nikolova *et al*.^51^ Signalling pathways were retrieved from KEGGREST (v1.44.1), and ligand genes, annotated in the CellChat^52^ ligand-receptor interaction database (CellChatDB; v2.2.0), were excluded from the expression-based pathway signatures to focus on downstream pathway activity. For each pathway, activity was quantified by combining expression of non-ligand pathway genes with Pando-derived transcription factor module activity. TF module activity was calculated using AUCell^74^ (v1.26.0), scoring positive and negative target modules into a single activity measure. Expression and module features were assembled into a cell-by-feature signature matrix, scaled across cells, and subjected to PCA. The first principal component was used as the pathway activity score, with its sign flipped when features were negatively correlated with the component. Scores were added to cell metadata and summarized by cell type and gestational age.

Intercellular communication was inferred using CellChat^52^ (v2.2.0) on the transcriptomic data. The truncated mean algorithm was used to identify signalling pathways, communication probabilities, and ligand-receptor pairs with interactions retained at *P* < 0.01. Cellular communication and ligand-receptor pairs were visualized using default CellChat plotting functions. For spatial CellChat analysis, the default distance parameter (10 µm) was enabled, with contact-dependent signalling disabled; spatial interactions were considered significant at *P* < 0.05.

---

**Figure S1:**
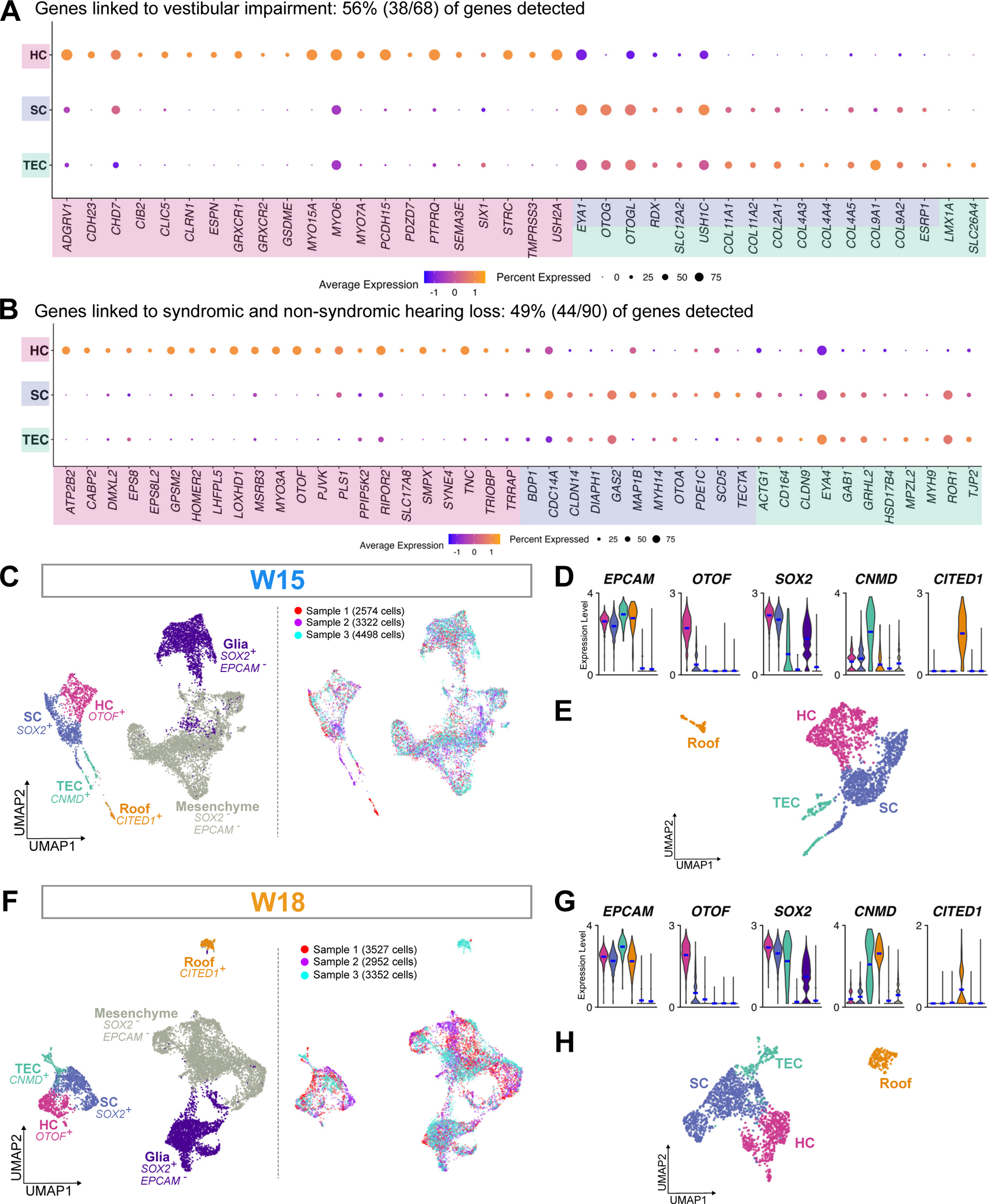
Genes involved in inner ear dysfunction expressed in the fetal utricle and spatial transcriptomic annotations; related to Figure 1. (A) Dot plot showing the genes associated with vestibular dysfunction in the integrated utricle dataset (n=2 per stage). Genes are detected in ≥25% of cells within a cell type. (B) Dot plot showing the genes associated with syndromic and non-syndromic hearing loss (n=2 per stage). (C) UMAP projection of cells from W15 Xenium cross-sections and sample distribution, n=3. (D) Violin plots of marker genes used for cell type annotation in the W15 Xenium dataset. (E) Re-clustered UMAP projection of W15 *EPCAM^+^* epithelial cells. (F) UMAP projection of cells from W18 Xenium cross-sections and sample distribution, n=3. (G) Violin plots of marker genes used for cell type annotation in the W18 Xenium dataset. (H) Re-clustered UMAP projection of W18 *EPCAM^+^* epithelial cells.

**Figure S2:**
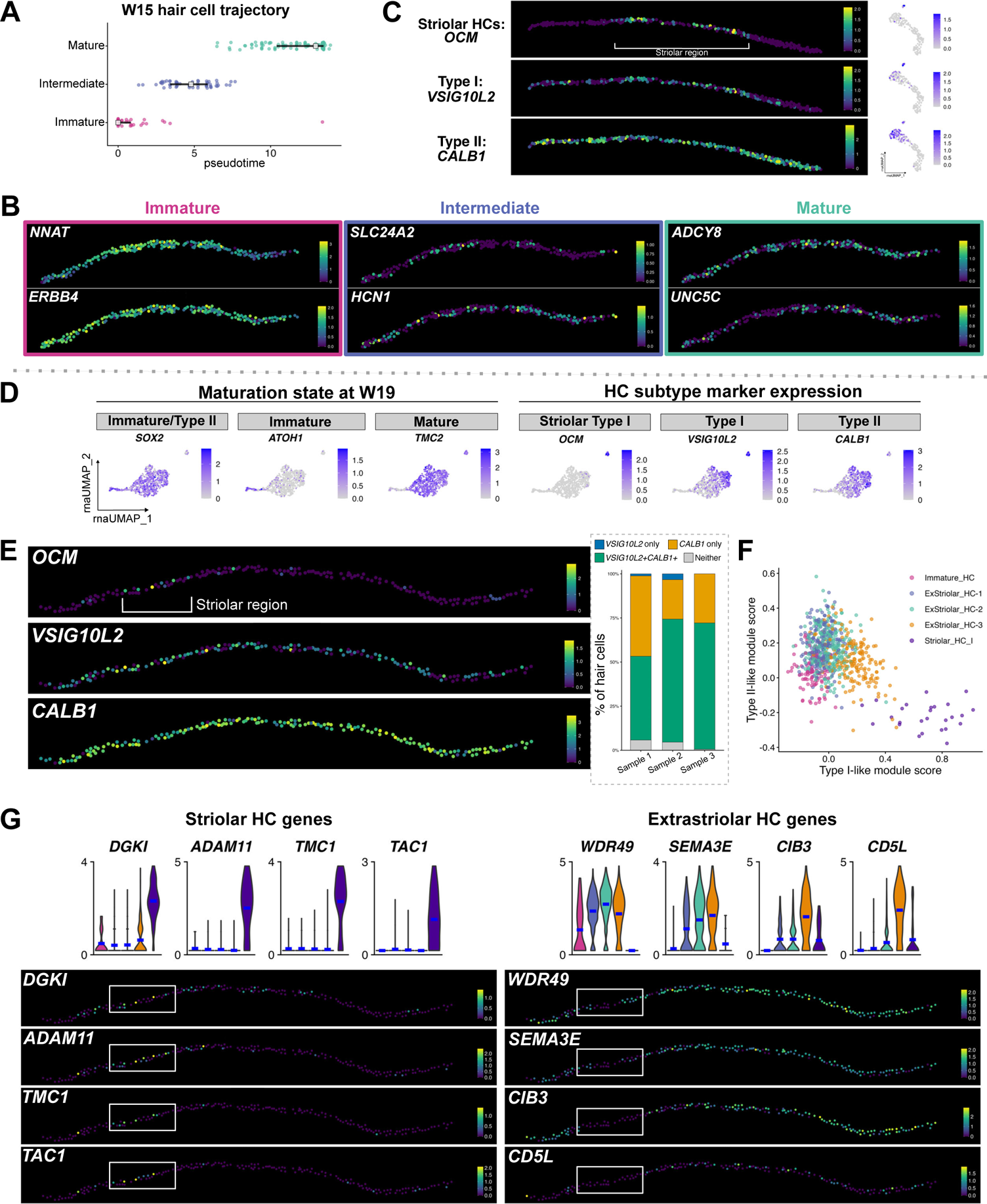
W15 and W19 hair cell identities; related to Figure 2. (A) Monocle3 pseudotime inference of W15 hair cells. White squares indicate median pseudotime value of each cluster and black lines indicate interquartile range. (B) Spatial feature plots of immature, intermediate, and mature hair cell cluster enriched genes (n=3). (C) Spatial expression (n=3) and feature plots of *OCM*, *VSIG10L2*, and *CALB1* in W15 hair cells. (D) Expression of *SOX2*, *ATOH1*, and *TMC2* as markers of hair cell maturation, as well as *OCM*, *VSIG10L2*, and *CALB1* for hair cell subtype specification. (E) Spatial feature plots showing distribution of *OCM*, *VSIG10L2,* and *CALB1* in W18 hair cells (n=3). Inset depicts the fraction of hair cells expressing *VSIG10L2* only*, CALB1* only, both markers, or neither marker. (F) Scatter plot of type I and type II hair cell module scores across W19 hair cell clusters. (G) Violin and spatial feature plots of region-specific hair cell genes.

**Figure S3:**
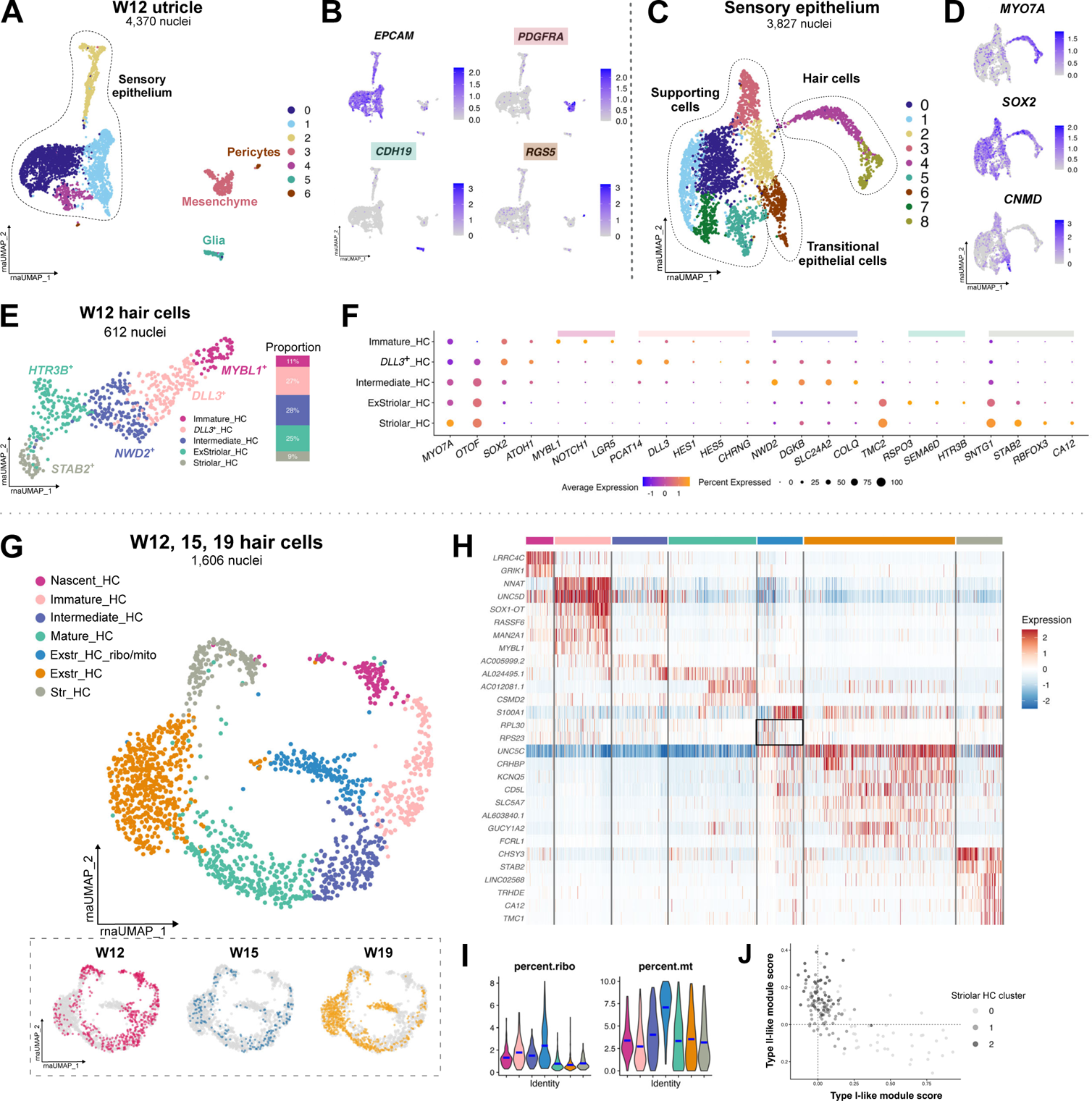
Cell types in the W12 utricle and supplementary hair cell integration; related to Figure 3. (A) UMAP projection of nuclei in the W12 utricle (n=1) based on snRNA-seq data. (B) Feature plots of cell type markers *EPCAM* (epithelial; clusters 0, 1, 2,4), *CDH19* (glia; cluster 5), *PDGFRA* (mesenchyme; cluster 3), and *RGS5* (pericytes; cluster 6). (C) Reclustering of *EPCAM^+^* cells from (A). (D) Feature plots of *MYO7A*, *SOX2*, and *CNMD* within *EPCAM^+^* cells from (C). (E) UMAP projection and cluster proportions of W12 hair cell clusters, n=1. (F) Dot plot of the top differentially expressed genes in each hair cell subpopulation. (G) UMAP projection of integrated hair cells from W12 to W19 prior to subsetting. Inset depicts distribution of cells from each age. (H) Heatmap of top differentially expressed genes in each cluster in (G). One extrastriolar cluster is marked by enriched expression of ribosomal genes (boxed). (I) Violin plots of mitochondrial and ribosomal genes across hair cell clusters. (J) Scatter plot of type I and type II hair cell module scores across striolar hair cell subclusters.

**Figure S4:**
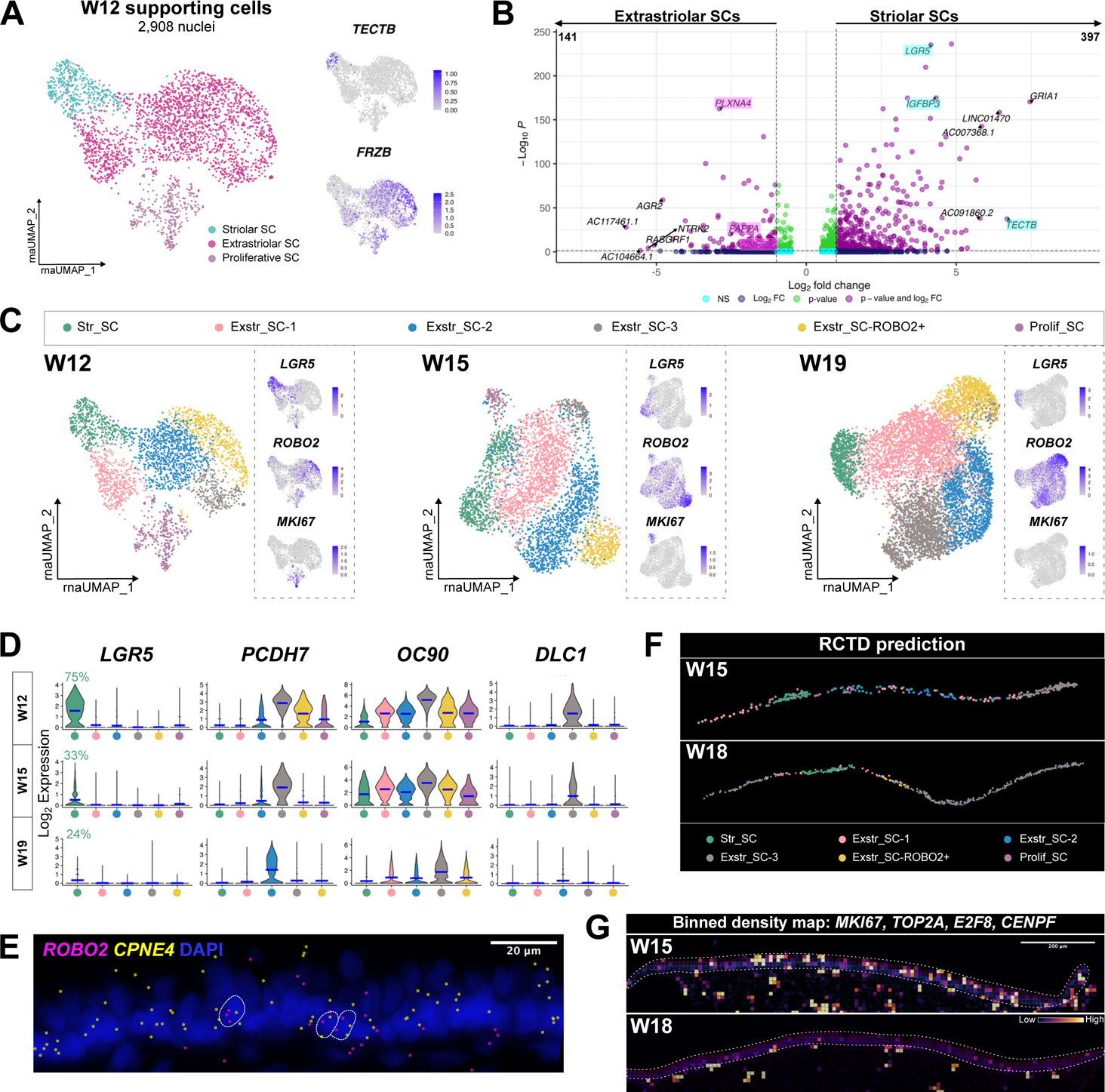
Supporting cell information; related to Figure 5. (A) W12 striolar and extrastriolar supporting cells. Inset shows feature plots of striolar marker *TECTB* and extrastriolar marker *FRZB*. (B) Volcano plot depicting differentially expressed genes between W12 striolar and extrastriolar supporting cells. Highlighted genes correspond to genes in Figure 5F. (C) Unsupervised clustering of supporting cells at W12, W15, and W19. Insets show feature plots of *LGR5*, *ROBO2*, and *MKI67*. (D) Violin plots of genes enriched in subclusters of supporting cells at each age. The percentage of *LGR5*^+^ striolar supporting cells are marked. (E) *In situ* expression of *ROBO2* in supporting cells (*CPNE4^+^*), n=3. (F) Predicted spatial identity of each supporting cell subcluster at W15 and W18 using Robust Cell Type Decomposition (RCTD), n=3. (G) Binned (10 µm) density maps of proliferation genes *MKI67*, *TOP2A*, *E2F8*, and *CENPF* in W15 and W18 utricles (n=3). Sensory epithelium is delineated with a white dashed line.

**Figure S5:**
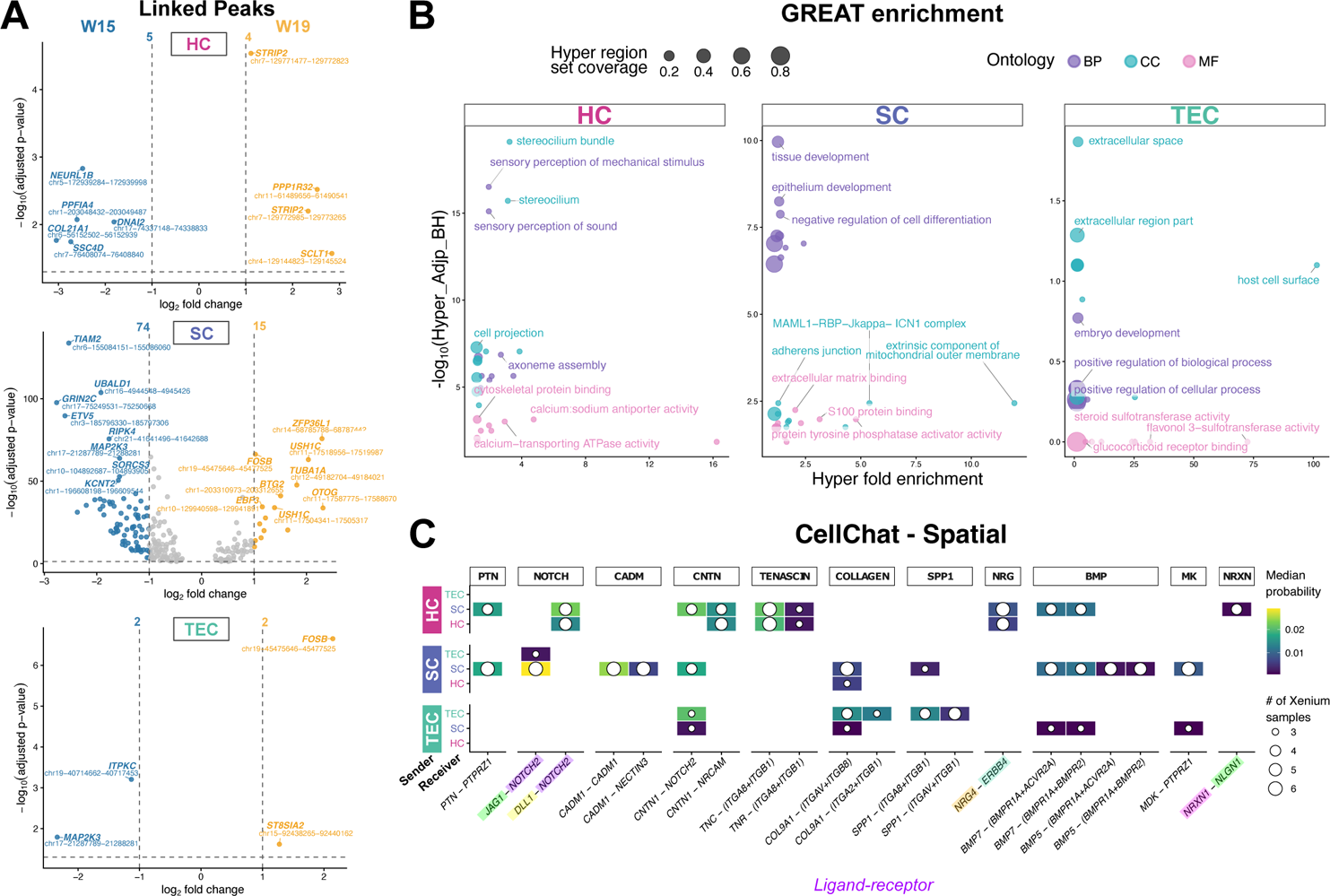
Epigenomic profiles of each cell type and CellChat signalling. Related to Figure 7. (A) Volcano plots of differentially accessible linked peaks between W15 and W19 within each cell type. (B) GREAT BP, CC, and MF terms for each cell type, thresholded at hyper fold enrichment ≥ 1 for visibility. The top three terms are labelled. (C) CellChat-inferred ligand-receptor (L-R) pairs accounting for spatial proximity in the Xenium dataset. LR pairs validated in Figure 7 (K-M) are highlighted.

## References

1. Rauch, S.D., Velazquez-Villaseñor, L., Dimitri, P.S., and Merchant, S.N. (2001). Decreasing Hair Cell Counts in Aging Humans. Annals of the New York Academy of Sciences 942, 220–227. 10.1111/j.1749-6632.2001.tb03748.x.

2. Merchant, S.N., Veläzquez-Villasenor, L., Tsuji, K., Glynn, R.J., Wall 3rd, C., and Rauch, S.D. (2000). Temporal Bone Studies of the Human Peripheral Vestibular System. 1. Normative Vestibular Hair Cell Data. Ann Otol Rhinol Laryngol Suppl . 109, 3–13. 10.1177/00034894001090s502.

3. Tsuji, K., Velazquez-Villaseñor, L., Rauch, S.D., Glynn, R.J., Wall 3rd, C., and Merchant, S.N. (2000). Temporal Bone Studies of the Human Peripheral Vestibular System. 3. Aminoglycoside Ototoxicity. Ann Otol Rhinol Laryngol Suppl . 109, 20–25. 10.1177/00034894001090S504.

4. Ding, D., Jiang, H., Zhang, J., Xu, X., Qi, W., Shi, H., Yin, S., and Salvi, R. (2018). Cisplatin-induced vestibular hair cell lesion-less damage at high doses. Journal of Otology 13, 115–121. 10.1016/j.joto.2018.08.002.

5. Fancello, V., Palma, S., Monzani, D., Pelucchi, S., Genovese, E., and Ciorba, A. (2021). Vertigo and Dizziness in Children: An Update. Children 8, 1025. 10.3390/children8111025.

6. Cao, C., Cade, W.T., Li, S., McMillan, J., Friedenreich, C., and Yang, L. (2021). Association of Balance Function With All-Cause and Cause-Specific Mortality Among US Adults. JAMA Otolaryngol Head Neck Surg 147, 460. 10.1001/jamaoto.2021.0057.

7. Agrawal, Y., Carey, J.P., Della Santina, C.C., Schubert, M.C., and Minor, L.B. (2009). Disorders of Balance and Vestibular Function in US Adults: Data From the National Health and Nutrition Examination Survey, 2001-2004. Arch Intern Med 169, 938. 10.1001/archinternmed.2009.66.

8. Jönsson, R., Sixt, E., Landahl, S., and Rosenhall, U. (2004). Prevalence of dizziness and vertigo in an urban elderly population. VES 14, 47–52. 10.3233/VES-2004-14105.

9. Taylor, R.R., Filia, A., Paredes, U., Asai, Y., Holt, J.R., Lovett, M., and Forge, A. (2018). Regenerating hair cells in vestibular sensory epithelia from humans. eLife 7, e34817. 10.7554/eLife.34817.

10. Wang, T., Ling, A.H., Billings, S.E., Hosseini, D.K., Vaisbuch, Y., Kim, G.S., Atkinson, P.J., Sayyid, Z.N., Aaron, K.A., Wagh, D., et al. (2024). Single-cell transcriptomic atlas reveals increased regeneration in diseased human inner ear balance organs. Nat Commun 15, 4833. 10.1038/s41467-024-48491-y.

11. Kim, G.S., Wang, T., Sayyid, Z.N., Fuhriman, J., Jones, S.M., and Cheng, A.G. (2022). Repair of surviving hair cells in the damaged mouse utricle. Proc. Natl. Acad. Sci. U.S.A. 119, e2116973119. 10.1073/pnas.2116973119.

12. Kawamoto, K., Izumikawa, M., Beyer, L.A., Atkin, G.M., and Raphael, Y. (2009). Spontaneous hair cell regeneration in the mouse utricle following gentamicin ototoxicity. Hearing Research 247, 17–26. 10.1016/j.heares.2008.08.010.

13. Wan, G., Corfas, G., and Stone, J.S. (2013). Inner ear supporting cells: Rethinking the silent majority. Seminars in Cell & Developmental Biology 24, 448–459. 10.1016/j.semcdb.2013.03.009.

14. Monzack, E.L., and Cunningham, L.L. (2013). Lead roles for supporting actors: Critical functions of inner ear supporting cells. Hearing Research 303, 20–29. 10.1016/j.heares.2013.01.008.

15. Burns, J.C., On, D., Baker, W., Collado, M.S., and Corwin, J.T. (2012). Over Half the Hair Cells in the Mouse Utricle First Appear After Birth, with Significant Numbers Originating from Early Postnatal Mitotic Production in Peripheral and Striolar Growth Zones. JARO 13, 609–627. 10.1007/s10162-012-0337-0.

16. Severinsen, S.Å., Sørensen, M.S., Kirkegaard, M., and Nyengaard, J.R. (2010). Stereological estimation of total cell numbers in the young human utricular macula. Acta Oto-Laryngologica 130, 773–779. 10.3109/00016480903397694.

17. Rosenhall, U., and Engström, B. (1974). Surface Structures of the Human Vestibular Sensory Regions. Acta Oto-Laryngologica 77, 3–18. 10.1080/16512251.1974.11675749.

18. Luca, E., Ibeh, N., Yamamoto, R., Liang, W., Bennett, D., Lin, V., Chen, J., Lovett, M., and Dabdoub, A. (2025). Revealing heterogeneity and damage response in the adult human utricle. Nat Commun 17, 9. 10.1038/s41467-025-66358-8.

19. Van Der Valk, W.H., Van Beelen, E.S.A., Steinhart, M.R., Nist-Lund, C., Osorio, D., De Groot, J.C.M.J., Sun, L., Van Benthem, P.P.G., Koehler, K.R., and Locher, H. (2023). A single-cell level comparison of human inner ear organoids with the human cochlea and vestibular organs. Cell Reports 42, 112623. 10.1016/j.celrep.2023.112623.

20. Hao, Y., Stuart, T., Kowalski, M.H., Choudhary, S., Hoffman, P., Hartman, A., Srivastava, A., Molla, G., Madad, S., Fernandez-Granda, C., et al. (2024). Dictionary learning for integrative, multimodal and scalable single-cell analysis. Nature Biotechnology 42, 293–304. 10.1038/s41587-023-01767-y.

21. Warchol, M.E., Massoodnia, R., Pujol, R., Cox, B.C., and Stone, J.S. (2019). Development of hair cell phenotype and calyx nerve terminals in the neonatal mouse utricle. J of Comparative Neurology 527, 1913–1928. 10.1002/cne.24658.

22. Hume, C.R., Bratt, D.L., and Oesterle, E.C. (2007). Expression of LHX3 and SOX2 during mouse inner ear development. Gene Expression Patterns 7, 798–807. 10.1016/j.modgep.2007.05.002.

23. Trapnell, C., Cacchiarelli, D., Grimsby, J., Pokharel, P., Li, S., Morse, M., Lennon, N.J., Livak, K.J., Mikkelsen, T.S., and Rinn, J.L. (2014). The dynamics and regulators of cell fate decisions are revealed by pseudotemporal ordering of single cells. Nat Biotechnol 32, 381–386. 10.1038/nbt.2859.

24. Yamada, S., Furukawa, R., and Sakakibara, S. (2022). Identification and expression profile of novel STAND gene Nwd2 in the mouse central nervous system. Gene Expression Patterns 46, 119284. 10.1016/j.gep.2022.119284.

25. Scheffer, D., Sage, C., Plazas, P.V., Huang, M., Wedemeyer, C., Zhang, D., Chen, Z., Elgoyhen, A.B., Corey, D.P., and Pingault, V. (2007). The α1 subunit of nicotinic acetylcholine receptors in the inner ear: transcriptional regulation by ATOH1 and co-expression with the γ subunit in hair cells. Journal of Neurochemistry 103, 2651–2664. 10.1111/j.1471-4159.2007.04980.x.

26. Poppi, L.A., Holt, J.C., Lim, R., and Brichta, A.M. (2020). A review of efferent cholinergic synaptic transmission in the vestibular periphery and its functional implications. Journal of Neurophysiology 123, 608–629. 10.1152/jn.00053.2019.

27. Holt, J.C., Kewin, K., Jordan, P.M., Cameron, P., Klapczynski, M., McIntosh, J.M., Crooks, P.A., Dwoskin, L.P., and Lysakowski, A. (2015). Pharmacologically Distinct Nicotinic Acetylcholine Receptors Drive Efferent-Mediated Excitation in Calyx-Bearing Vestibular Afferents. J. Neurosci. 35, 3625–3643. 10.1523/JNEUROSCI.3388-14.2015.

28. Burns, J.C., Kelly, M.C., Hoa, M., Morell, R.J., and Kelley, M.W. (2015). Single-cell RNA-Seq resolves cellular complexity in sensory organs from the neonatal inner ear. Nat Commun 6, 8557. 10.1038/ncomms9557.

29. Simmons, D.D., Tong, B., Schrader, A.D., and Hornak, A.J. (2010). Oncomodulin identifies different hair cell types in the mammalian inner ear. J of Comparative Neurology 518, 3785–3802. 10.1002/cne.22424.

30. Wang, X., Liu, S., Cheng, Q., Qu, C., Ren, R., Du, H., Li, N., Yan, K., Wang, Y., Xiong, W., et al. (2023). CIB2 and CIB3 Regulate Stereocilia Maintenance and Mechanoelectrical Transduction in Mouse Vestibular Hair Cells. J. Neurosci. 43, 3219–3231. 10.1523/JNEUROSCI.1807-22.2023.

31. Ratzan, E.M., Lee, J., Madison, M.A., Zhu, H., Zhou, W., Géléoc, G.S.G., and Holt, J.R. (2024). TMC function, dysfunction, and restoration in mouse vestibular organs. Front. Neurol. 15, 1356614. 10.3389/fneur.2024.1356614.

32. Takimoto, Y., Ishida, Y., Nakamura, Y., Kamakura, T., Yamada, T., Kondo, M., Kitahara, T., Uno, A., Imai, T., Horii, A., et al. (2014). 5-HT3 receptor expression in the mouse vestibular ganglion. Brain Research 1557, 74–82. 10.1016/j.brainres.2014.02.016.

33. Hartman, B.H., Hayashi, T., Nelson, B.R., Bermingham-McDonogh, O., and Reh, T.A. (2007). Dll3 is expressed in developing hair cells in the mammalian cochlea. Developmental Dynamics 236, 2875–2883. 10.1002/dvdy.21307.

34. Ni, C., Wei, Y., Vona, B., Park, D., Wei, Y., Schmitz, D.A., Ding, Y., Sakurai, M., Ballard, E., Li, L., et al. (2025). A programmed decline in ribosome levels governs human early neurodevelopment. Nat Cell Biol 27, 1240–1255. 10.1038/s41556-025-01708-8.

35. McLean, C.Y., Bristor, D., Hiller, M., Clarke, S.L., Schaar, B.T., Lowe, C.B., Wenger, A.M., and Bejerano, G. (2010). GREAT improves functional interpretation of cis-regulatory regions. Nat Biotechnol 28, 495–501. 10.1038/nbt.1630.

36. Fleck, J.S., Jansen, S.M.J., Wollny, D., Zenk, F., Seimiya, M., Jain, A., Okamoto, R., Santel, M., He, Z., Camp, J.G., et al. (2023). Inferring and perturbing cell fate regulomes in human brain organoids. Nature 621, 365–372. 10.1038/s41586-022-05279-8.

37. Huang, E.J., Liu, W., Fritzsch, B., Bianchi, L.M., Reichardt, L.F., and Xiang, M. (2001). Brn3a is a transcriptional regulator of soma size, target field innervation and axon pathfinding of inner ear sensory neurons. Development 128, 2421–2432. 10.1242/dev.128.13.2421.

38. Shi, T., Beaulieu, M.O., Saunders, L.M., Fabian, P., Trapnell, C., Segil, N., Crump, J.G., and Raible, D.W. (2023). Single-cell transcriptomic profiling of the zebrafish inner ear reveals molecularly distinct hair cell and supporting cell subtypes. eLife 12, e82978. 10.7554/eLife.82978.

39. Caspa Gokulan, R., Yap, L.F., and Paterson, I.C. (2022). HOPX: A Unique Homeodomain Protein in Development and Tumor Suppression. Cancers 14, 2764. 10.3390/cancers14112764.

40. Ono, K., Keller, J., López Ramírez, O., González Garrido, A., Zobeiri, O.A., Chang, H.H.V., Vijayakumar, S., Ayiotis, A., Duester, G., Della Santina, C.C., et al. (2020). Retinoic acid degradation shapes zonal development of vestibular organs and sensitivity to transient linear accelerations. Nat Commun 11, 63. 10.1038/s41467-019-13710-4.

41. Wang, S., Ibrahim, L.A., Kim, Y.J., Gibson, D.A., Leung, H.C., Yuan, W., Zhang, K.K., Tao, H.W., Ma, L., and Zhang, L.I. (2013). Slit/Robo Signaling Mediates Spatial Positioning of Spiral Ganglion Neurons during Development of Cochlear Innervation. Journal of Neuroscience 33, 12242–12254. 10.1523/JNEUROSCI.5736-12.2013.

42. Cable, D.M., Murray, E., Zou, L.S., Goeva, A., Macosko, E.Z., Chen, F., and Irizarry, R.A. (2022). Robust decomposition of cell type mixtures in spatial transcriptomics. Nat Biotechnol 40, 517–526. 10.1038/s41587-021-00830-w.

43. Zhao, X., Yang, H., Yamoah, E.N., and Lundberg, Y.W. (2007). Gene targeting reveals the role of Oc90 as the essential organizer of the otoconial organic matrix. Developmental Biology 304, 508–524. 10.1016/j.ydbio.2007.01.013.

44. Burns, J.C., Cox, B.C., Thiede, B.R., Zuo, J., and Corwin, J.T. (2012). *In Vivo* Proliferative Regeneration of Balance Hair Cells in Newborn Mice. J. Neurosci. 32, 6570–6577. 10.1523/JNEUROSCI.6274-11.2012.

45. Ishii, M., Tateya, T., Matsuda, M., and Hirashima, T. (2021). Stalling interkinetic nuclear migration in curved pseudostratified epithelium of developing cochlea. R. Soc. open sci. 8, 211024. 10.1098/rsos.211024.

46. Jan, T.A., Eltawil, Y., Ling, A.H., Chen, L., Ellwanger, D.C., Heller, S., and Cheng, A.G. (2021). Spatiotemporal dynamics of inner ear sensory and non-sensory cells revealed by single-cell transcriptomics. Cell Reports 36, 109358. 10.1016/j.celrep.2021.109358.

47. Chen, Z., Żak, M., Xu, S., De Andrés, J., and Daudet, N. (2025). A tissue boundary orchestrates the segregation of inner ear sensory organs. 10.1101/2022.03.03.482809.

48. Watanabe, K., and Ogawa, A. (1984). Carbonic Anhydrase Activity in Stria Vascularis and Dark Cells in Vestibular Labyrinth. Ann Otol Rhinol Laryngol 93, 262–266. 10.1177/000348948409300315.

49. Rayamajhi, D., Ege, M., Ukhanov, K., Ringers, C., Zhang, Y., Jung, I., D’Gama, P.P., Li, S.S., Cosacak, M.I., Kizil, C., et al. (2024). The forkhead transcription factor Foxj1 controls vertebrate olfactory cilia biogenesis and sensory neuron differentiation. PLoS Biol 22, e3002468. 10.1371/journal.pbio.3002468.

50. Elkon, R., Milon, B., Morrison, L., Shah, M., Vijayakumar, S., Racherla, M., Leitch, C.C., Silipino, L., Hadi, S., Weiss-Gayet, M., et al. (2015). RFX transcription factors are essential for hearing in mice. Nat Commun 6, 8549. 10.1038/ncomms9549.

51. Nikolova, M.T., He, Z., Seimiya, M., Jonsson, G., Cao, W., Okuda, R., Wimmer, R.A., Okamoto, R., Nikoloff, J.M., Dittrich, P.S., et al. (2025). Fate and state transitions during human blood vessel organoid development. Cell 188, 3329–3348.e31. 10.1016/j.cell.2025.03.037.

52. Jin, S., Guerrero-Juarez, C.F., Zhang, L., Chang, I., Ramos, R., Kuan, C.-H., Myung, P., Plikus, M.V., and Nie, Q. (2021). Inference and analysis of cell-cell communication using CellChat. Nat Commun 12, 1088. 10.1038/s41467-021-21246-9.

53. David, A.P., Biswas, S., Soltis, M.P., Eltawil, Y., Zhou, R., Easow, S.A., Cheng, A.G., Heller, S., and Jan, T.A. (2025). Crosstalk Signaling Between the Epithelial and Non-Epithelial Compartments of the Mouse Inner Ear. JARO 26, 127–145. 10.1007/s10162-025-00980-7.

54. You, D., Guo, J., Zhang, Y., Guo, L., Lu, X., Huang, X., Sun, S., and Li, H. (2022). The heterogeneity of mammalian utricular cells over the course of development. Clinical & Translational Med 12, e1052. 10.1002/ctm2.1052.

55. Wang, T., Chai, R., Kim, G.S., Pham, N., Jansson, L., Nguyen, D.-H., Kuo, B., May, L.A., Zuo, J., Cunningham, L.L., et al. (2015). Lgr5+ cells regenerate hair cells via proliferation and direct transdifferentiation in damaged neonatal mouse utricle. Nat Commun 6, 6613. 10.1038/ncomms7613.

56. Lin, J., Zhang, X., Wu, F., and Lin, W. (2015). Hair cell damage recruited Lgr5-expressing cells are hair cell progenitors in neonatal mouse utricle. Front. Cell. Neurosci. 9. 10.3389/fncel.2015.00113.

57. Shu, Y., Li, W., Huang, M., Quan, Y.-Z., Scheffer, D., Tian, C., Tao, Y., Liu, X., Hochedlinger, K., Indzhykulian, A.A., et al. (2019). Renewed proliferation in adult mouse cochlea and regeneration of hair cells. Nat Commun 10, 5530. 10.1038/s41467-019-13157-7.

58. Jen, H.-I., Hill, M.C., Tao, L., Sheng, K., Cao, W., Zhang, H., Yu, H.V., Llamas, J., Zong, C., Martin, J.F., et al. (2019). Transcriptomic and epigenetic regulation of hair cell regeneration in the mouse utricle and its potentiation by Atoh1. eLife 8, e44328. 10.7554/eLife.44328.

59. Roccio, M., and Edge, A.S.B. (2019). Inner ear organoids: new tools to understand neurosensory cell development, degeneration and regeneration. Development 146, dev177188. 10.1242/dev.177188.

60. Bucks, S.A., Cox, B.C., Vlosich, B.A., Manning, J.P., Nguyen, T.B., and Stone, J.S. (2017). Supporting cells remove and replace sensory receptor hair cells in a balance organ of adult mice. eLife 6, e18128. 10.7554/eLife.18128.

61. González-Garrido, A., Pujol, R., López-Ramírez, O., Finkbeiner, C., Eatock, R.A., and Stone, J.S. (2021). The Differentiation Status of Hair Cells That Regenerate Naturally in the Vestibular Inner Ear of the Adult Mouse. J. Neurosci. 41, 7779–7796. 10.1523/JNEUROSCI.3127-20.2021.

62. Wang, T., Niwa, M., Sayyid, Z.N., Hosseini, D.K., Pham, N., Jones, S.M., Ricci, A.J., and Cheng, A.G. (2019). Uncoordinated maturation of developing and regenerating postnatal mammalian vestibular hair cells. PLoS Biol 17, e3000326. 10.1371/journal.pbio.3000326.

63. Sayyid, Z.N., Wang, T., Chen, L., Jones, S.M., and Cheng, A.G. (2019). Atoh1 Directs Regeneration and Functional Recovery of the Mature Mouse Vestibular System. Cell Reports 28, 312–324.e4. 10.1016/j.celrep.2019.06.028.

64. Hern, W.M. (1984). Correlation of Fetal Age and Measurements Between 10 and 26 Weeks of Gestation. Obstetrics Gynecol. 63, 26–32.

65. Deng, Y., Ehiogu, B., Luca, E., Dabdoub, A., Lê Cao, K.-A., Wells, C.A., and Nayagam, B.A. (2025). Trophic and temporal dynamics of macrophage biology in human inner ear organogenesis. Front. Immunol. 16, 1690583. 10.3389/fimmu.2025.1690583.

66. Young, M.D., and Behjati, S. (2020). SoupX removes ambient RNA contamination from droplet-based single-cell RNA sequencing data. GigaScience 9, giaa151. 10.1093/gigascience/giaa151.

67. Hafemeister, C., and Satija, R. (2019). Normalization and variance stabilization of single-cell RNA-seq data using regularized negative binomial regression. Genome Biol 20, 296. 10.1186/s13059-019-1874-1.

68. Zappia, L., and Oshlack, A. (2018). Clustering trees: a visualization for evaluating clusterings at multiple resolutions. GigaScience 7, giy083. 10.1093/gigascience/giy083.

69. Gaspar, J.M. (2018). Improved peak-calling with MACS2. Preprint at Bioinformatics, 10.1101/496521 https://doi.org/10.1101/496521.

70. Stuart, T., Srivastava, A., Madad, S., Lareau, C.A., and Satija, R. (2021). Single-cell chromatin state analysis with Signac. Nature Methods 18, 1333–1341. 10.1038/s41592-021-01282-5.

71. Wang, T., Ling, A.H., Billings, S.E., Hosseini, D.K., Vaisbuch, Y., Kim, G.S., Atkinson, P.J., Sayyid, Z.N., Aaron, K.A., Wagh, D., et al. (2024). Single-cell transcriptomic atlas reveals increased regeneration in diseased human inner ear balance organs. Nat Commun 15, 4833. 10.1038/s41467-024-48491-y.

72. Yu, G., Wang, L.-G., Han, Y., and He, Q.-Y. (2012). clusterProfiler: an R Package for Comparing Biological Themes Among Gene Clusters. OMICS: A Journal of Integrative Biology 16, 284–287. 10.1089/omi.2011.0118.

73. Gu, Z., and Hübschmann, D. (2023). rGREAT: an R/bioconductor package for functional enrichment on genomic regions. Bioinformatics 39, btac745. 10.1093/bioinformatics/btac745.

74. Aibar, S., González-Blas, C.B., Moerman, T., Huynh-Thu, V.A., Imrichova, H., Hulselmans, G., Rambow, F., Marine, J.-C., Geurts, P., Aerts, J., et al. (2017). SCENIC: single-cell regulatory network inference and clustering. Nat Methods 14, 1083–1086. 10.1038/nmeth.4463.Figure 1: Cell types of the human fetal utricle.

